# Mechanism and function of root circumnutation

**DOI:** 10.1101/2020.05.04.075127

**Authors:** Isaiah Taylor, Kevin Lehner, Erin McCaskey, Niba Nirmal, Yasemin Ozkan-Aydin, Mason Murray-Cooper, Rashmi Jain, Elliot W. Hawkes, Pamela C. Ronald, Daniel I. Goldman, Philip N. Benfey

## Abstract

Early root growth is critical for plant establishment and survival. We have identified a molecular pathway required for oscillatory root tip movement known as circumnutation. Here we report a multiscale investigation of the regulation and function of this phenomenon. We identify key cell signaling events comprising interaction of the ethylene, cytokinin, and auxin hormone signaling pathways. We identify the gene *Oryza sativa Histidine Kinase-1/OsHK1*, as well as the auxin influx carrier gene *OsAUX1*, as essential regulators of this process in rice. Robophysical modelling demonstrated the benefits of tip movement for navigating past obstacles, prompting us to challenge mutant and wild-type plants with different substrates. Consistent with model behavior, root circumnutation facilitated exploration of a solid surface and promoted seedling establishment in rocky soil. Thus, the integration of robotics, physics and biology elucidated the functional importance of root circumnutation and uncovered the molecular mechanisms underlying its regulation.

**One sentence summary:** Circumnutation facilitates root exploration.

---

During the life cycle of a plant, penetration of roots into soil is a fundamental yet physically challenging task (*1*, *2*). This is especially true for the primary root (the first root to emerge from the seed), because it provides the seedling with its sole source of anchorage and water/nutrient absorption, and because it must enter the soil absent other mechanical support (*3*, *4*). The emergent primary root faces two related challenges: it must grow into a heterogeneous soil substrate wherever the seed lands, and it must do so when the seed is otherwise unanchored. Species adapted to growing in wetlands, such as rice, face the added complication of establishing during sporadic floods that pose a catastrophic danger of washout (*5*–*7*). Thus, the earliest stages of root growth represent a critical window of survival when rapid and robust root penetration into soil is essential. It is therefore logical to posit that primary root growth strategies have evolved to meet the challenges arising from the myriad growth conditions seedling roots encounter.

## Results

To identify regulators of such early root growth strategies in rice, we performed a genetic screen on a subset of a fully sequenced mutant population in the model rice cultivar KitaakeX (*8*–*10*). Plants were grown in transparent gel media, and the developing root systems were imaged over the course of several days using a previously described phenotyping system (*11*, *12*). From this screen, we isolated 3 allelic mutants each exhibiting an approximately 50% increase in primary root length, all harboring lesions in the *Oryza sativa HISTIDINE KINASE-1/HK1* gene (Fig. 1A, Fig. S1). To understand the dynamics of root elongation in the mutants, we used a modified version of our phenotyping platform employing automated time-lapse imaging to monitor growth every 15 minutes for several days following germination. This imaging revealed wild-type roots undergoing striking oscillatory movement of the tip known as circumnutation, with the mutants appearing to grow predominantly downward (Movie S1). To visualize this phenomenon, we tracked the position of the root tip in wild type and the mutants for two days after germination. We then estimated a central axis of root growth using locally estimated scatterplot smoothing (loess) regression, and plotted displacement from this central axis at each 15-minute time point (Fig. 1B, Fig. S1). From this analysis we observed that in wild type, circumnutation typically becomes regular within a day of germination and approaches a maximum in both amplitude and period by approximately 48 hours. In contrast, the mutant never enters into an organized circumnutational pattern and the root tip remains situated close to the central axis of growth.

**Figure 1).**
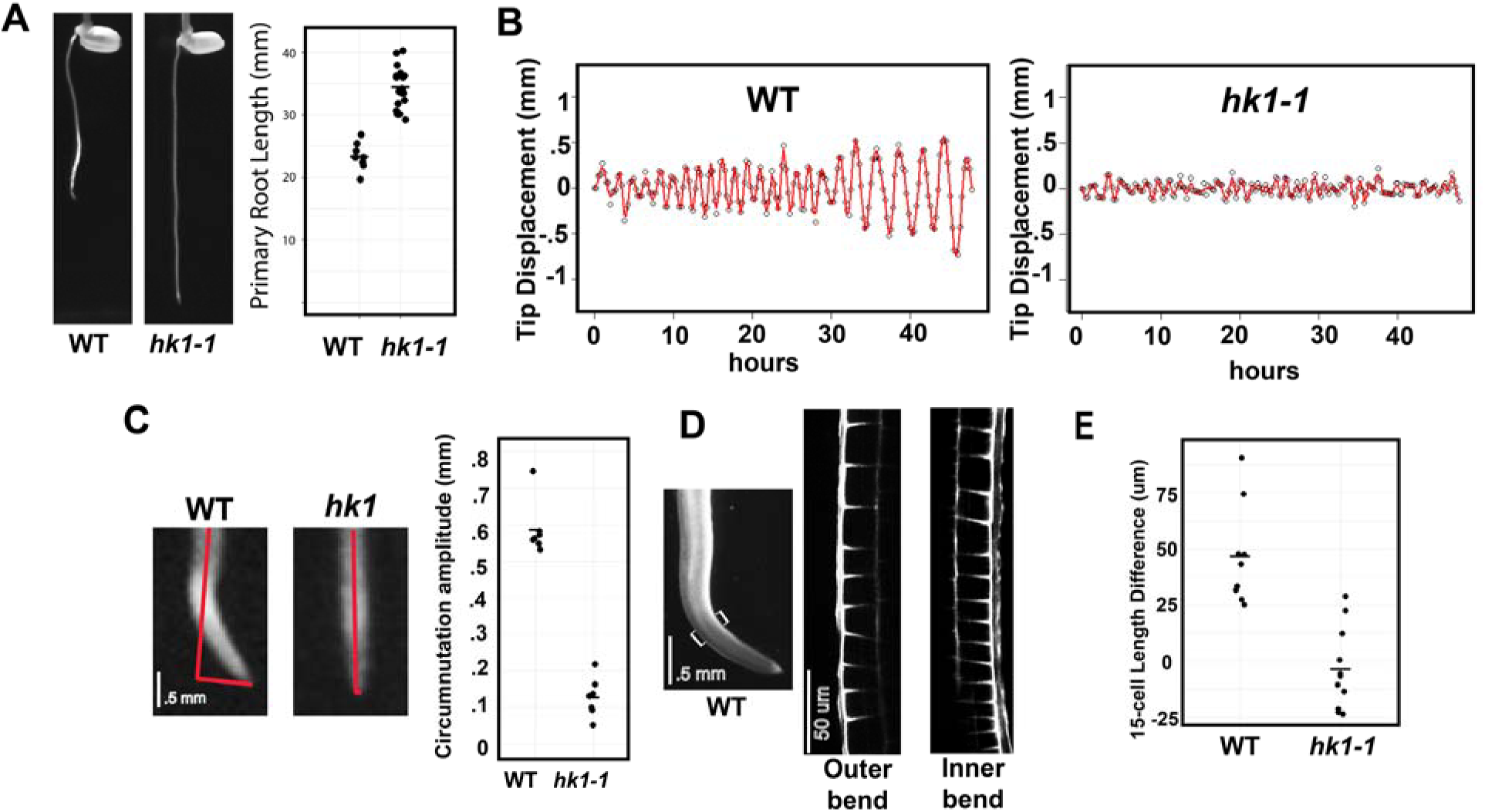
Phenotypic analysis of the *histidine kinase-1/hk1* mutant. A) Primary root depth 2 days after germination for wild type and the *hk1-1* mutant. N = 8 and 18 primary roots for wild type and mutant, respectively. 2-sided Wilcoxon Rank Sum test p-value = 7.1e-05. Horizontal dash indicates mean value. B) Distance of root tip from central axis of growth every 15 minutes for 48 hours after germination. Red plots represent natural cubic splines fit to the data for visualization. C) Quantification of average radius of circumnutation in wild type and *hk1* between 40-48 hours after germination. N = 7 primary roots for each genotype. 2-sided Wilcoxon Rank Sum test p-value = 0.0005828. Horizontal dash indicates mean value. D) Example circumnutating wild-type primary root exhibiting higher epidermal cell elongation on the outer bend compared to the inner bend. E) Quantification of the difference between outer and inner length of the region of 15 epidermal cells flanking the maximal bend in wild type and *hk1-1* mutant control. N = 9 and 11 primary roots for wild type and mutant, respectively. 2-sided Wilcoxon Signed Rank test p-values = 0.003906 and 0.5771 for wild type and mutant, respectively. Horizontal dash indicates mean value.

In wild-type roots, circumnutation is driven by a pronounced bending of the root, typically occurring 1-2 mm from the tip, in the region of rapid cellular elongation (*13*). To quantify this phenotype, we calculated the amplitude of tip displacement for several wild-type roots over the course of 8 hours after circumnutation had been strongly initiated (Fig. 1C, Materials and Methods). In this experiment, wild type had a mean amplitude of tip displacement equaling 0.60 mm (±.07 mm SD), which corresponds to the radius of the cylinder circumscribed by the root tip as it revolves around its central axis (Fig. 1C). In contrast, the mutant exhibits erratic, small amplitude displacement with a mean of approximately 0.13 mm (±.05 mm SD) over the same timeframe (Wilcoxon Rank Sum test p-value = 0.0005828, Fig. 1C).

Root bending during circumnutation is reminiscent of root bending during response to gravitational stimulus, a process driven by reduced cell elongation on the lower side of the turning root (reviewed in (*14*)). Examination of epidermal cells on the outer and inner bend of circumnutating wild-type roots revealed a consistent decrease in elongation on the inner bend (Fig. 1D), suggesting there is similar mechanical regulation of root bending occurring during circumnutation. To quantify this effect, we recorded the epidermal cell length on the inner and outer side of 9 circumnutating wild-type roots in a region of 15 cells flanking a line bisecting the root at the region of maximal bending (Fig. S2). We calculated the difference between the length of this region of cells on the outer and inner side of the root and found a consistent decrease in 15-cell length on the inner side of the bend (Wilcoxon Signed Rank test p-value = 0.003906, Fig. 1E). As a control, we examined 11 *hk1-1* mutant roots. Because mutant roots exhibit no pronounced bending, we defined bisecting lines at comparable distances from the root tip and randomly labelled the sides of the root. The distribution of length differences in the mutant was centered around 0, with no root exhibiting strong bias for length on one side or the other bend (Wilcoxon Signed Rank test p-value = 0.5771, Fig. 1E). Combined with the difference in primary root length between wild type and mutant, this result suggests circumnutation is driven by transient, circumferentially localized restriction of cell elongation on the inner side of the bending root.

The occurrence of root circumnutation in rice has been described in a number of studies (*15*–*17*), and evidence for root circumnutation has been observed in other species since Charles and Francis Darwin’s work on plant tropisms (*18*). However, the mechanistic regulation and function of this process are still largely undefined. *HK1* has recently been shown to positively regulate ethylene signaling in rice roots downstream of ethylene receptors (*19*). This signaling pathway was shown to restrict root elongation, but a connection between ethylene signaling and circumnutation has not been established. To assess if ethylene signaling regulates circumnutation, we applied the ethylene receptor inhibitor 1-methylcyclopropene (1-MCP) to circumnutating wild-type roots. This treatment led to a rapid and dramatic cessation of circumnutation and a sudden increase in the rate of root elongation (Fig. S3, Movie S2). This result is consistent with a model where ethylene activates HK1 to promote circumnutation, leading to a reduction in root elongation.

We next focused on understanding the mechanism of downstream regulation of circumnutation. *HK1* encodes an active histidine kinase with homology to cytokinin receptors, but lacking a predicted cytokinin binding domain (*19*, *20*) (Fig. S4). Ethylene has been shown to mediate activation of the HK1 protein leading to direct phosphorylation of histidine-containing phosphotransfer/AHP proteins, well-characterized effectors that function downstream of receptors in canonical cytokinin signaling (*19*). This HK1 mediated AHP phosphorylation in turn feeds into cytokinin responsive transcriptional regulation (*19*). These results indicate regulation of *HK1* represents a previously unobserved point of cross-talk between ethylene and cytokinin signaling.

We reasoned if the *HK1* pathway activates downstream cytokinin signaling, and reduction of this signaling causes the *hk1* mutant phenotype, we could rescue circumnutation by treating the mutant with exogenous cytokinin. Consistent with this hypothesis, treatment with the naturally occurring cytokinin, *trans*-zeatin, led to a rescue of circumnutation and an increase in the mean amplitude of tip displacement from 0.11 mm (±.03 mm SD) in untreated control roots to 0.64 mm (±.11 mm SD) in *trans*-zeatin treated roots (Wilcoxon Rank Sum test p-value = 0.01587, Fig. 2A, Movie S3). RNA-Seq analysis of the distal 2mm of root tip revealed that transcript abundance levels of canonical cytokinin signaling-induced type A response regulator genes are reduced between 45%-90% in the *hk1* mutant (Fig. 2B, Data S1). Similar RNA-seq analysis demonstrated that this reduction is reversed upon treatment with *trans*-zeatin (Fig. 2B). Furthermore, we found a strong, positive relationship between global gene expression changes comparing log(fold change) of wild type/mutant with log(fold change) of mutant plus/minus *trans*-zeatin (Fig. S5). These results indicate HK1 activates downstream cytokinin signaling to regulate circumnutation. An initial diagram of our findings is displayed in Fig. 2C. An open question for future study is whether canonical cytokinin receptor signaling also makes a meaningful contribution to the regulation of circumnutation, or if *trans*-zeatin induced rescue of *hk1* represents artificial stimulation of downstream signaling by exogenous cytokinin.

**Figure 2).**
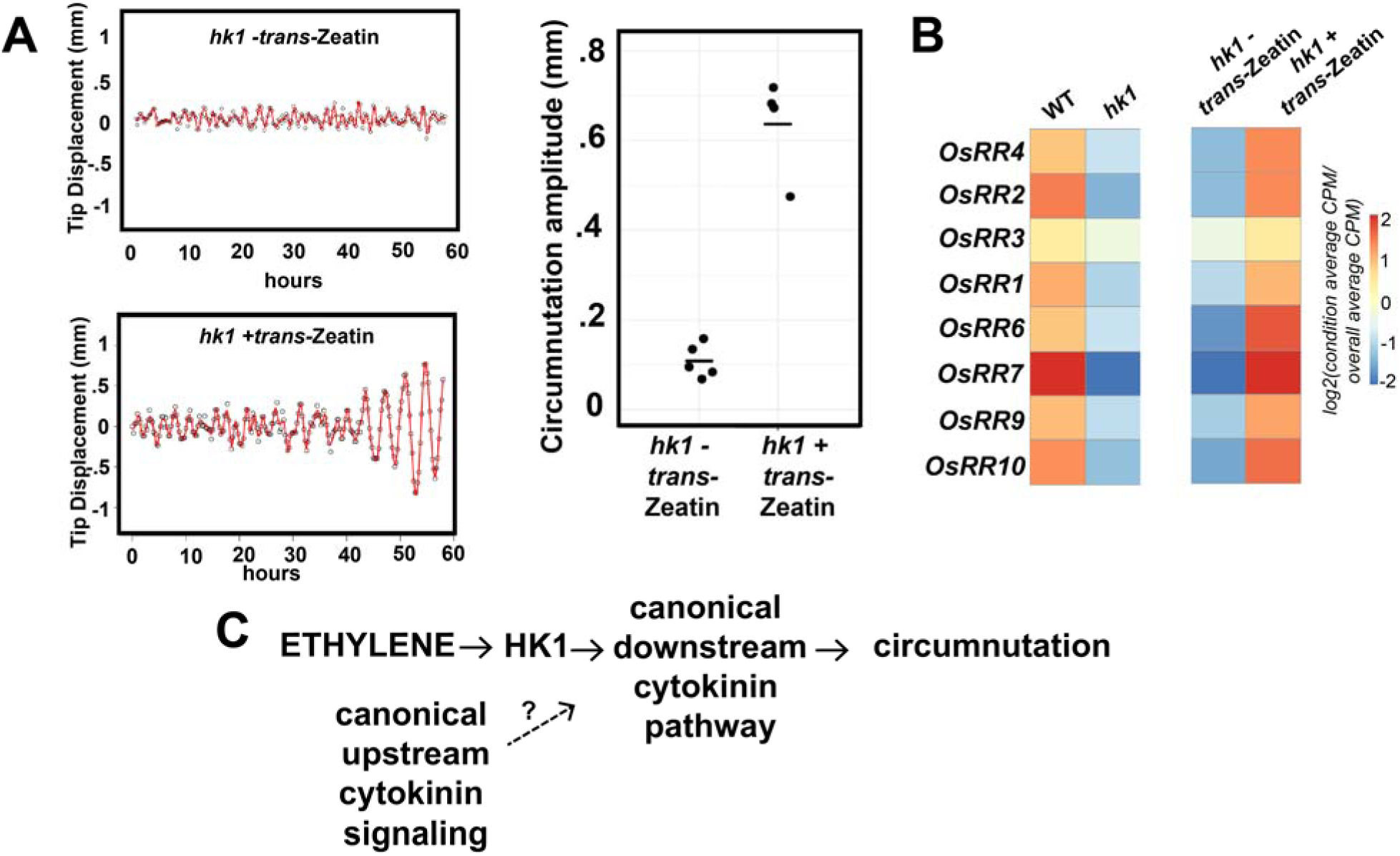
Characterization of molecular signaling downstream of *HK1*. A) Example root tip traces and quantification of average radius of circumnutation in *hk1* treated with *trans*-zeatin between 50-58 hours after germination. N = 5 and 4 primary roots for untreated and treated, respectively. 2-sided Wilcoxon Rank Sum test p-value = 0.01587. Horizontal dash indicates mean value. B) Relative expression level of 8 root tip expressed cytokinin signaling regulated Type-A Response Regulator genes in wild type vs. *hk1* and *hk1* +/− cytokinin. C) Diagram of proposed signaling pathway regulating of circumnutation. The uncertain contribution of canonical upstream cytokinin signaling to circumnutation is indicated by a question mark.

We next sought a mechanism to link this signaling pathway to regulation of cell elongation during circumnutation. Root bending during the graviresponse is driven by directional transport of the plant hormone auxin to the outer cell layers of the lower side of the root by the coordinated action of auxin influx and efflux carriers (reviewed in (*14*, *21*, *22*)). Accumulation of auxin on the lower side of the root then inhibits cell elongation, causing downward tip reorientation. Application of auxin transport inhibitors to the roots of pea seedlings has been shown to block circumnutation (*23*), suggesting a connection between auxin transport and root bending during circumnutation. Additionally, regulation of auxin transport and signaling have been implicated in a variety of oscillatory root growth phenomena in Arabidopsis (*24*–*31*), although interpretation of the underlying physical basis for these growth responses has been challenging (*28*, *32*, *33*).

To test the involvement of auxin in regulating circumnutation, we first examined our RNA-Seq data and discovered that in the *hk1* mutant there is a reduction in a large number of canonical auxin response genes such as those encoding AUX/IAA proteins (Fig. 3A) and *SMALL AUXIN UP-REGULATE*D*/SAUR* genes (Fig. S6). We also noted a general restoration of expression for these genes in the mutant treated with *trans*-zeatin (Fig. 3A, Fig. S6), consistent with the hypothesis that there is involvement of an auxin response in regulating circumnutation. To functionally test this hypothesis, we grew wild-type roots in media containing the auxin efflux inhibitor N-1-naphthylphthalamidic acid (NPA). This treatment resulted in near complete inhibition of circumnutation, as evidenced by a reduction in the mean amplitude of tip displacement from 0.62 mm (±.10 mm SD) in untreated control roots to 0.11 mm (±.12 mm SD) in NPA treated roots (Wilcoxon Rank Sum test p-value = 0.004329, Fig. 3B, Movie S4), indicating auxin transport likely plays a key role during this process. Next, we reasoned if the phenotype of *hk1* is due to defects in auxin transport, we may be able to rescue circumnutation in the mutant by addition of exogenous auxin. Because auxin has a potent inhibitory effect on root elongation, we allowed plants to grow untreated for approximately one day in hydroponic conditions before supplementing the media with auxin. We found that the predominant naturally occurring auxin, indole-3-acetic acid (IAA), did not rescue the circumnutation defect of *hk1* at any concentration tested (Movie S5). In contrast, the membrane permeable synthetic auxin 1-naphthaleneacetic acid (1-NAA), but not the structurally related non-auxinic analog 2-NAA, rescued the defect of the *hk1* mutant, with an increase in mean tip displacement from 0.19 mm (±.07 mm SD) in untreated control roots to 0.59 mm (±.14 mm SD) in roots treated with 1-NAA (Wilcoxon Rank Sum test p-value = 0.0005828, Fig. 3C, Movie S5).

**Figure 3).**
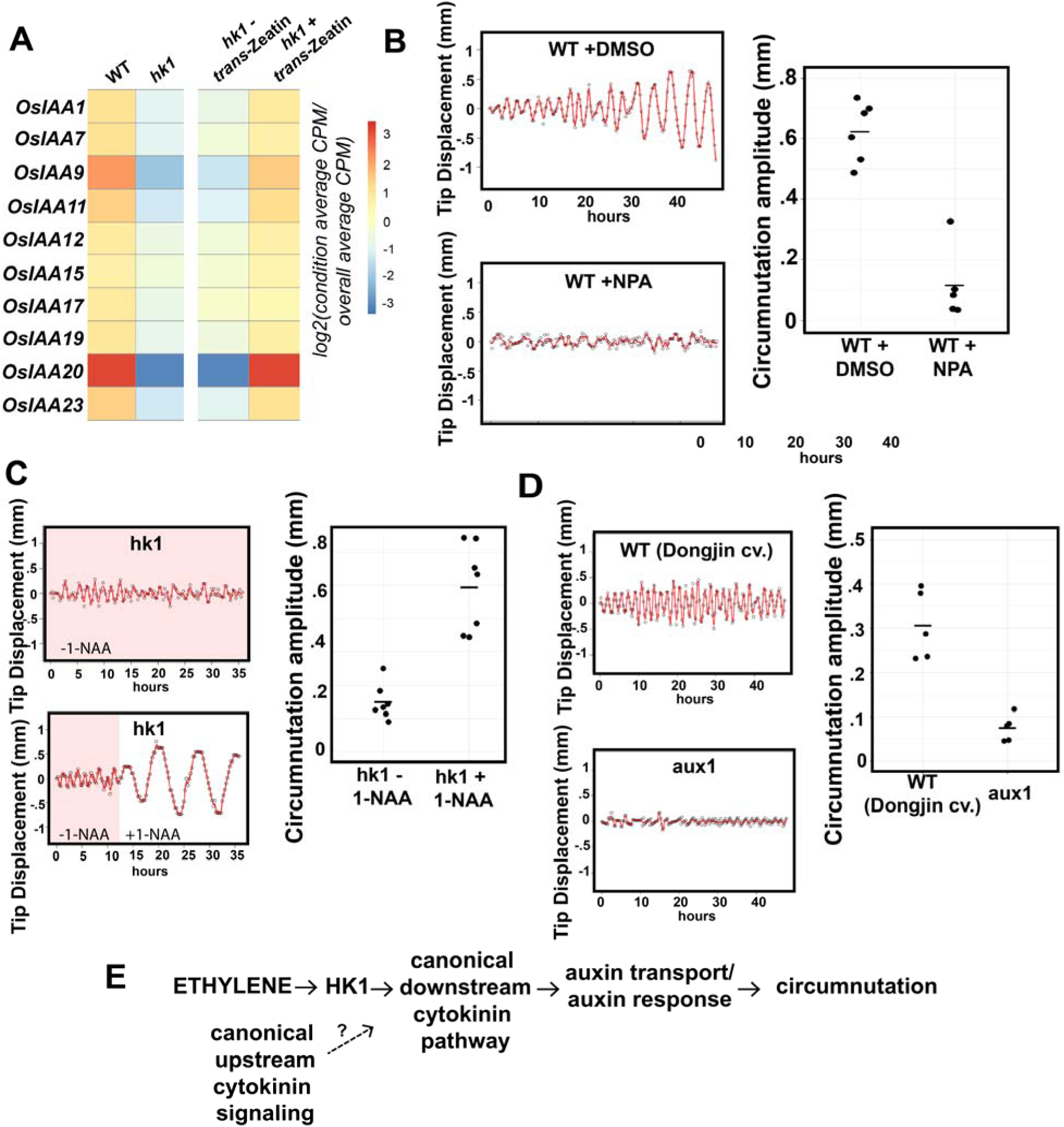
Evidence auxin transport and response regulate circumnutation. A) Relative expression level of 10 root tip expressed *AUX/IAA* genes in wild type vs. *hk1* and *hk1* +/− cytokinin. B) Example root tip traces and quantification of average radius of circumnutation in wild-type plants treated with NPA. N = 6 and 5 primary roots for untreated and treated, respectively. 2-sided Wilcoxon Rank Sum test p-value = 0.004329. Horizontal dash indicates mean value. C) Example root tip traces and quantification of average radius of circumnutation in *hk1* plants grown hydroponically after addition of 200 nM 1-NAA. N = 7 primary roots for both untreated and treated. 2-sided Wilcoxon Rank Sum test p-value = 0.0005828. Horizontal dash indicates mean value. D) Example root tip traces and quantification of average radius of circumnutation in wild type (Dongjin cultivar) and *aux1*. N = 5 primary roots for each genotype. 2-sided Wilcoxon Rank Sum test p-value = 0.007937. Horizontal dash indicates mean value. E) Diagram of proposed signaling pathway regulating of circumnutation. The uncertain contribution of canonical upstream cytokinin signaling to the regulation of circumnutation is indicated by a question mark.

The specific ability of 1-NAA to rescue circumnutation suggests a model of regulation by *HK1*. Because of its lipophilic structure, 1-NAA readily diffuses across the plasma membrane and does not require an auxin importer for rapid uptake from the apoplast, after which it is then an efficient substrate for the auxin efflux machinery (*34*). Thus, the ability of 1-NAA to rescue the *hk1* phenotype suggests auxin import, but not export, is critically impaired in the *hk1* mutant. To investigate this possibility, we re-examined our RNA-Seq data and noted transcript levels of the auxin importer gene *OsAUX1* are reduced approximately 30% in the *hk1* mutant, and its abundance is restored to approximately wild-type levels in the mutant treated with cytokinin (Fig. S7). To test if *AUX1* is a regulator of circumnutation, we analyzed an *aux1* mutant isolated in the Dongjin cultivar of rice and found, in addition to its previously described agravitropic and enhanced primary root length phenotypes (*35*, *36*), it exhibited a complete loss of circumnutation as evidenced by a reduction in the mean amplitude of tip displacement from 0.30 mm (±.08 mm SD) in wild type to .07 mm (±.03 mm SD) in *aux1* (Wilcoxon Rank Sum test p-value = 0.007937, Fig 3D, Movie S6). Collectively our results indicate HK1 regulates an auxin response controlling circumnutation, at least in part, by controlling AUX1-mediated auxin influx capacity. An updated diagram based on these findings is presented in Fig. 3E.

A previous study of root circumnutation across diverse rice cultivars established a positive correlation between root tip angle during circumnutation in laboratory experiments and the success of seedling establishment in field conditions (*15*), leading the authors to hypothesize that circumnutation promotes primary root colonization of soil. To directly test this hypothesis, we performed functional studies on wild type and the *hk1* mutant in artificial and natural growth substrates. The first experiment was motivated by a phenotype we observed when mutant roots reach the bottom of a growth container, where they often form restricted coils, whereas wild type grows along the surface distal to the site of contact (Fig. 4A). We hypothesized that this coiling behavior of *hk1* mutants would reduce its ability to effectively explore dense interfaces. To model this phenomenon, we used plastic surfaces with 2.5mm diameter holes equally spaced at different densities in a square lattice. The surfaces were placed on hollow platforms of equal height in containers with gel-based media and imaged with a high-throughput automated system (Fig. 4B). At the highest hole density (5mm spacing), both wild-type and *hk1* primary roots were effective in encountering holes and continuing deeper growth (Fig. 4C). However, as hole density decreased, *hk1* roots showed a pronounced reduction in success in encountering a hole (Fig. 2C, Movie S7). To quantify this effect, we employed logistic regression to model the probability of success in finding a hole as a function of spacing and genotype (Fig. S9). We observed an effect of genotype translating to an overall estimated 12.44-fold increase in the odds of success in the wild type compared to mutant (estimated odds ratio, 95% Wald CI (3.43, 45.09)). These data indicate that wild-type roots are more effective in exploration and less affected by sparse hole density than *hk1* mutants, providing a plausible mechanism to buffer against environmental uncertainty inherent in exploration of soil interfaces.

**Figure 4).**
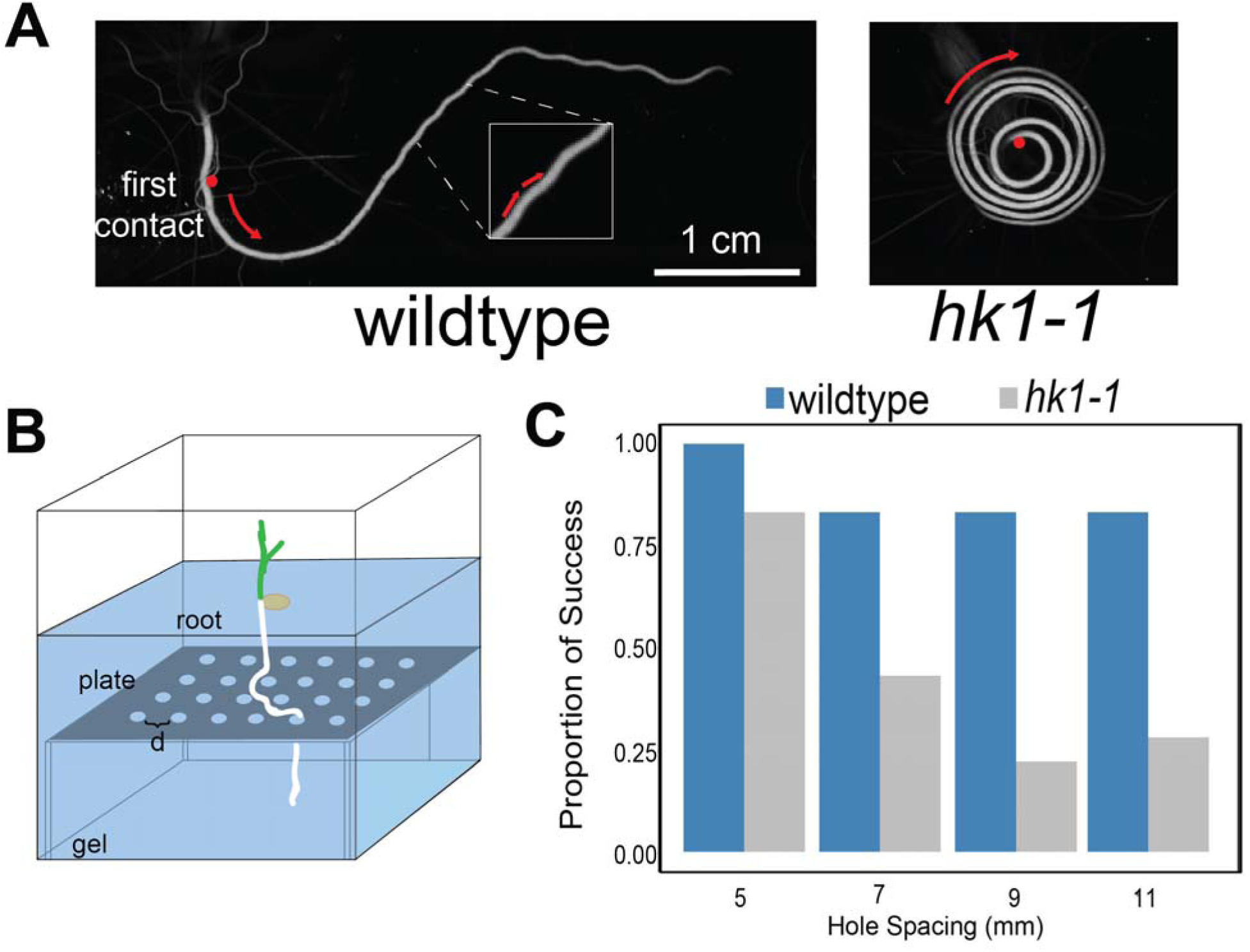
*hk1* has reduced surface exploration capacity. A) Example root growth after striking a transparent solid surface for wild-type and *hk1-1* mutant primary roots. Red dots indicates the point of first contact with the surface and red arrows represent direction of root growth. B) Schematic of experimental set-up to measure surface exploration. Platforms with holes separated by variable distances (d) were covered in gel-solidified media. Seeds were sown in the media and roots allowed to grow onto the platforms. C) Quantification of success in growing through a hole for wild-type and *hk1-1* mutant primary roots at different hole spacings. N = 11, 12, 6, and 6 roots for wild type and 12, 16, 18, and 7 for *hk1* at 5mm, 7mm, 9mm, and 11mm hole spacings, respectively.

To study mechanical principles underlying the function of circumnutation, we challenged a pneumatically actuated soft robophysical root model (described in (*37*, *38*)) with or without nutation in a variety of 2-dimensional obstacle courses (Movie S8). This device employs tip-based extension and is capable of generating oscillatory 2-dimensional nutations isolated to the distal terminus when side actuators are sequentially inflated (Fig. 5A, Fig. 5B). Two differences between this model and biological root growth are that the robot grows and oscillates in only two-dimensions, instead of three, and that the robotic root shortens the inner surface of a curve to pull the tip laterally while the biological root lengthens the outer surface to push the tip. Despite these differences, the model is useful by enabling dissection of the functional consequences of addition of a lateral growth component during root exploration in an experimentally tractable robotic system (*39*). Prior experiments demonstrated that nutating robotic roots are able to efficiently extend into a lattice of regularly spaced pegs, while non-nutating control robotic roots nearly always become stuck in the first two rows of the lattice^37^. To gain insight into the mechanism by which nutation improves penetration, we utilized video analysis of this experimental system to study what occurred when the robotic root encountered an obstacle. We hypothesized that nutating robotic roots would be able to contact obstacles nearly head-on yet pass, while non-nutating roots could only pass if they graze an obstacle. This is because nutation adds a lateral component to the force vector that the tip applies to the obstacle, making it more tangential; without nutation, this tip force is aligned with the axis of the root (Fig. 5D). Slipping along an obstacle to pass it only occurs when the tip force is sufficiently tangential to overcome the force of friction; thus a non-nutating root with an axis-aligned tip force must contact an obstacle more tangentially than a nutating root (Fig. 5D). To test this hypothesis, we measured the contact angle of the tip (α, see inset of Fig. 5E, Fig. S8) and compared the distribution of the contact angles where the robotic root successfully passed with and without nutation. In the nutating case, at even the lowest contact angles (most head-on contact), the success rate was over 80%. However, when the contact angle of a non-nutating root was less than 15°, the root tip always stuck to the peg and could never grow farther (Fig. 5C, Fig. S8). To derive a single measure of penetration effectiveness we calculated the proportion of times any root struck a peg and was able to continue growing (Fig. S8 and Materials and Methods). Overall, 97% of collisions of a nutating root with a peg resulted in a successful traversal, whereas only 36% of collisions by non-nutating roots resulted in a traversal (Chi-Squared test p-value = 7.768e-10, Fig. S8). These results indicate that a simple oscillatory motion may be sufficient for emergence of improved root exploratory ability.

**Figure 5).**
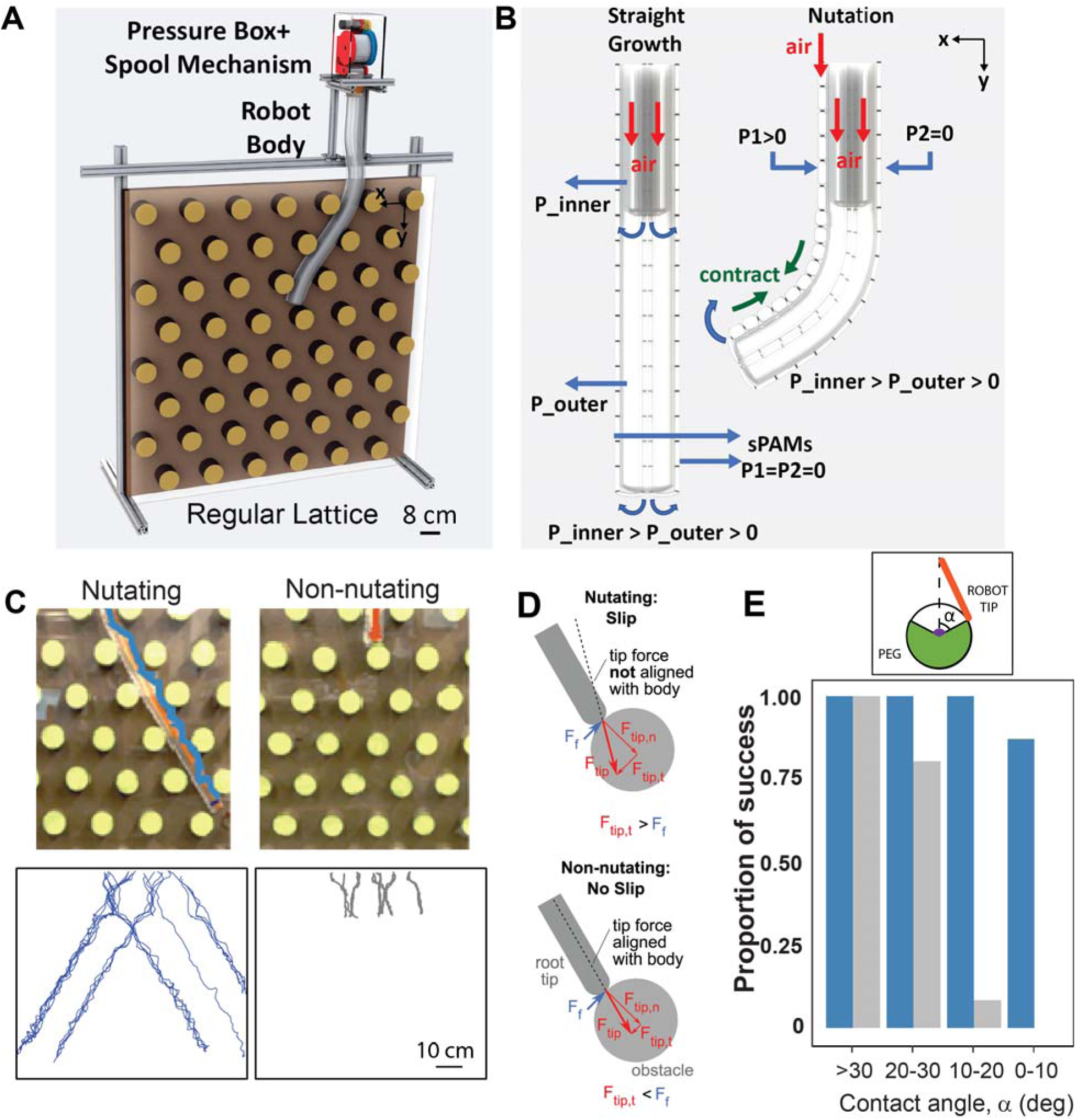
Robophysical model of root growth in heterogeneous substrates. A) Drawing of obstacle course and pneumatically actuated robotic root. The main body of the robot is attached to the pressure control box, and the spool mechanism simultaneously winds/unwinds the growing part of the robot during expansion/rewinding. B) When the pressure of the inner body is greater than the pressure of the outer tip body and both are greater than zero gauge, (P_inner>P_outer>0), the robot grows with tip eversion. Only the lower pressure tip bends if one of the side actuators (sPAMs) is activated (P1 or P2 > 0). C) (top) Representative images of the robot at the most common growth length with (left) and without (right) nutation. (bottom) Trajectories of nutating with a period of 5s (blue, 21 experiments) and non-nutating (gray, 21 experiments) robot roots through the regular lattice of round pegs starting from the different initial positions. D) For the non-nutating case, the tip force is aligned with the body; this means that the tangential component of the tip force (F_tip,t) is only created through misalignment of the tip with the obstacle. For nearly head-on collisions, F_tip,t will be less than the force of friction, F_f, and no slip occurs. For the nutating case, the tip force is not always aligned with the body, and the oscillating lateral force means that F_tip,t can exceed F_f to cause slip and passage of the obstacle. E) The proportion of obstacles that the robot tip successfully passed with nutation (blue, 21 experiments with a total of N = 17, 12, 15, and 15 contacts) and without nutation (gray, 21 experiments with N = 7, 5, 12, and 9 contacts) in each contact angle range (α). The inset shows the contact angle when the tip (orange) hits the peg (green).

**Figure 6).**
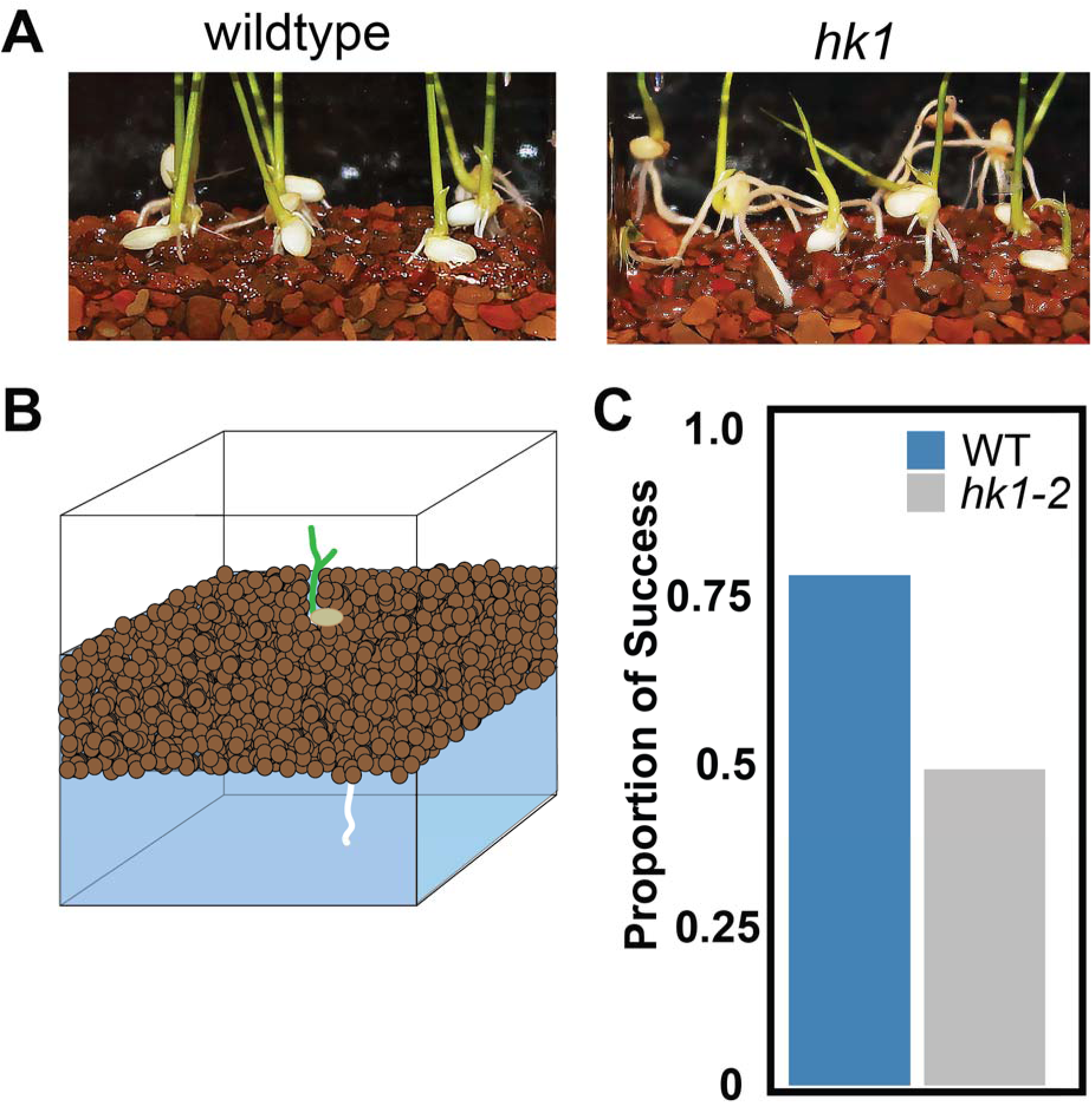
*hk1* is defective in penetration of granular substrates. A) Example root growth on liquid saturated gravel surface for wild-type and *hk1* mutant primary roots. B) Schematic of experimental set-up to measure gravel penetration. C) Quantification of success in growing through a 1.5 cm layer of gravel within 3 days of germination. N = 88 and 71 roots for wild type and *hk1-2*, respectively. 2-sided Chi-Squared test p-value = 0.0001253.

Our platform experiment and robotic simulation suggests root exploration can be understood as a probabilistic process where impenetrable obstacles along the growth path of the root represent potential failure points. We therefore hypothesized that circumnutation is an adaptation to facilitate root penetration in soil containing dense heterogeneities such as rocks or compaction. To test this hypothesis, we germinated wild-type and *hk1* mutant seeds on irregularly shaped calcined clay gravel of a defined size distribution saturated with liquid growth media. This experiment models the situation encountered by the seeds of semi-aquatic plant species such as rice adapted to germination in heterogeneous, water-logged soil conditions. We noted a striking phenomenon where the roots of wild type would effectively burrow into the gravel layer, while mutant seedlings’ roots often became caught near the surface, leading to dislodging of the seeds, rolling of the shoot downward, and aerial exposure of long root segments in protruding arcs (Fig. 5A, Movie S9). We quantified this effect by planting seeds of wild type and mutant on 1.5 cm of gravel overlaying a gel layer and counting the number of primary roots emerging into the gel layer 3 days post-germination (Fig. 5B, Movie S10). In this experiment we observed 77% of wild-type roots penetrating into the gel layer, but only 48% of mutant roots (Chi-Squared test p-value = 0.000238, Fig. 5C). Thus, our robotic modelling of circumnutation was accurate in predicting the reduction in penetration ability of mutant roots on a heterogeneous growth substrate, and indicates the simulation, despite its simplicity, captures an essential mechanical function of circumnutation. Collectively these results indicate root circumnutation promotes seedling establishment by aiding in effective exploration of dense, heterogeneous substrates.

Our results provide support for a model in which *HK1* signaling positively regulates establishment of a mobile signal transported cell to cell in a concerted tangential direction, negatively regulating cell elongation, producing circumnutation. Our genetic and physiological results suggest that the signal is auxin, given that circumnutation appears to require both auxin influx and efflux, and auxin is known to both be directionally transported and to negatively regulate root cellular elongation. Once circumnutation has been initiated, it appears to be employed by the root as a robust exploratory strategy. This is likely to be of critical importance to primary roots in natural growth settings, which emerge when the seed is unanchored. Searching for a path of least resistance into the soil is potentially a preferable strategy to attempting concerted downward growth. This highlights a key conclusion from our study, that there appears to be a tradeoff between rapid, unidirectional root elongation and multidimensional root exploration. We anticipate continued study of circumnutation will provide additional insight into regulation of exploratory root growth.

## Acknowledgments

This work was supported by a grant from the NSF (NSF PHY-1915445) to DIG and PNB, by a NSF Graduate Research Fellowship to KRL, by the Howard Hughes Medical Institute and the Gordon and Betty Moore Foundation (through Grant GBMF3405) to PNB, by the Dunn Family Professorship to DIG and ENM, U.S. National Science Foundation (Award #1237975), the Foundation for Food and Agricultural Research (Award # 534683) and the National Institutes of Health (GM122968) to PCR. The authors would like to thank Medhavinee Mijar, Carrie Hunner, Milan Shah, Enes Aydin, and Anupam Mijar for assistance in collecting and analyzing data, the staff of the Duke Phytotron for plant care, Duke GCB for Illumina sequencing services, and Drs. Lucia Strader, Sabrina Sabbatini, Jazz Dickinson, Trevor Nolan, Edith Pierre-Jerome, and Pablo Szekely for critical reading of the manuscript, as well as Drs. Joe Kieber and Flor Ercoli for helpful discussions.

## Author contributions

Conceptualization: IWT, KRL, DIG, PNB. Methodology: IWT, KRL, ENM, NN, YOA, MMC, RJ, PCR. Investigation : IWT, KRL, ENM, NN, YOA, MMC, RJ, PCR. Data Curation: RJ. Writing – original draft preparation: IWT, KRL, ENM, YOA. Writing – review and editing: all authors. Supervision: EWH, PCR, DIG, PNB.

## Data and materials availability

All data is available in the main text, the supplementary materials, or in public repositories listed in the text.

## Materials and Methods

### Plant Materials and Growth Conditions

The *hk1* mutants are in the KitaakeX background, a wildtype Kitaake line transformed with the *XA21* gene driven by the maize (Zea mays) ubiquitin promoter (*10*, *40*). *oshk1-1* is from the mutant line FN-287 and has a SBS at position Chr6:26808461. *oshk1-2* is from FN-179 and has an inversion between Chr6:25948672-26809852. *oshk1-3* is from FN790-S and has a deletion from Chr6:26807814-26812817. Genomic coordinates are from Os-Nipponbare-Reference-IRGSP-1.0 (*41*). The *osaux1-3* mutant is in the Dongjin cultivar and contains a T-DNA insertion in exon 6 (*42*).

Seeds were dehulled, sterilized with 50% bleach for 10 minutes, and rinsed 4-5 times with sterile water. All plant growth was in Yoshida’s nutrient solution solidified with gellan gum (Gelzan, Caisson Inc.) (*43*).

For time-lapse imaging experiments, sterilized seeds were grown directly in GA-7 Magenta vessels containing media solidified with 0.15% Gelzan, except auxin treatment experiments where plants were grown in sterilized, cut pipet tip racks supported by fiberglass mesh floating in liquid Yoshida’s medium, and the surface exploration experiment where plants were grown in .5% gelzan solidified Yoshida’s media. After planting, seeds were incubated for two days in the dark at 30°C, vessels were then moved to constant, low light (approximately 650 lux) at 22-23°C for imaging, except the surface exploration experiment which was conducted under 12 hour light/dark cycles with night-time illumination when the camera imaged each shelf.

### Time-lapse Imaging

We designed and constructed two robotic automatic imaging systems to acquire images of rice plants growing in Magenta containers at 15-minute intervals. The robot were capable of simultaneously imaging 100 and 85 containers, respectively. FLIR Flea3 cameras (FL3-U3-13S2C-CS or FL3-U3-13S2M-CS) were positioned facing the front of the growth containers to visualize root tip circumnutation and root depth. The cameras were moved at set intervals by Arduino-controlled horizontal and vertical gantries (UPC 819368022278, OpenBuilds) with NEMA 17 stepper motors (UPC 819368021264 and 819368021226, OpenBuids). Illumination was provided by LED striplights (Novostella, B075RYSHQQ).

### Analysis of Primary Root Length

The length of the primary roots from a sample of wildtype and *hk1-1, hk-2,* and *hk1-3* were calculated by measuring the roots from photos of the respective genotypes two days post-germination. Germination was defined as the point when the primary root had emerged and grown to a depth equal to the bottom of the seed. An analysis script, along with the raw data, has been included in Data S1.

### Statistical Analysis of Primary Root Length

We performed 2-sided Wilcoxon Rank Sum tests comparing wildtype and mutant primary root lengths. A script performing this procedure, along with raw data, has been included in Data S1.

### Root Tip Tracing

Root tip tracing was performed by marking the position of the root tip in successive images in ImageJ. These coordinates were input into R and an estimated central axis of root growth was calculated using loess regression. We then calaculated the distance of the root tip from this central axis of root growth at each timepoint and fit a highly flexible natural cubic spline model using the splines package in R in order to visualize the path of the root tip. Conversion from pixel to distance was accomplished by calculating a conversion factor derived from an image of a standard metric ruler placed at the same distance from the camera as the roots typically grow. A script performing these functions, along with raw data, has been included in Data S2.

### Circumnutation Amplitude Quantification

For quantification of circumnutation amplitude for wildtype KitaakeX vs *hk1,* wildtype KitaakeX +/− 1-MCP, *hk1* +/− cytokinin, wildtype KitaakeX +/− NPA, and wildtype Dongjin vs. *aux1, w*e modified the above script to identify local extrema in the measurements of root tip displacement within an 8 hour window after circumnutation had increased to a maximum amplitude in the given condition. In wildtype KitaakeX and Dongjin, we found circumnutation was at or near maximum amplitude 40-48 hours after germination, and in cytokinin treated *hk1* mutant we found it occurred between 50-58 hours after germination. We then averaged the displacement of the root tip from the central axis of growth at each local extrema during this time window to calculate an average circumnutation amplitude and compared it to the negative control over the same timeframe. In the 1-MCP experiment, we allowed roots to grow for approximately 2 days and then treated with 1-MCP. We measured circumnutation amplitude in the time period 2-10 hours after addition. A similar procedure was performed for *hk1* +/− 1-NAA, except due to the extreme reduction in root elongation rate observed after 1-NAA treatment, we instead averaged the 4 final local extrema observed during the approximately 24 hours of imaging with the hormone (two maxima and two minima).

### Statistical Analysis of Circumnutation Amplitude

We performed 2-sided Wilcoxon Rank Sum tests comparing control and test condition circumnutation amplitudes in each respective experiment. A script performing this procedure, along with raw data, has been included in Data S3.

### Root Cell Length Measurements

Roots of 2-3 day old seedlings were assayed for cell length measurements. In wildtype, circumnutating roots were defined as those with at least 15 degree bending of the root tip. Roots were live-stained with 1mg/ml calcofluor white or 10 mg/ml propidium iodide for 5 minutes and immediately imaged using a Zeiss 510 or Zeiss 880 confocal microscope. A combined tile scan/z-stack of the root tip and elongation zone of the roots was used to capture overall root shape and cell structure. In ImageJ, a line perpendicular to the direction of root growth was used to mark the region of the root exhibiting maximal bending. We then measured the longitudinal cell length at the region of maximal bending for 15 epidermal cells on the inner and outer flanks of the root (9 cells rootward, 5 cells shootward, and 1 cell at the position of maximum bending). We then took the difference of the total length of the 15 cells on the outer and inner bend, the result of which is plotted in Fig. 1E. For the mutant, we performed the same procedure, making a line at approximately 1 mm from the root tip and measuring cells on both sides of the root before arbitrarily subtracting the length of the measured cells from one side of the root from the length of the cells on the other.

### Statistical Analysis of Root Cell Length

We performed a 2-sided Wilcoxon Signed Rank test comparing the difference in 15-cell length from one side of the root to the other under the null hypothesis that they are centered on 0. A script performing this procedure, along with raw data, has been included in Data S4.

### Hormone Treatment/Inhibitor Experiments

For the 1-MCP experiment, we grew seeds of wildtype in magenta boxes, as above, and approximately 2 days after germination when circumnutation had been strongly initiated, we added freshly prepared .014% 1-MCP mixed with 1 ml of water sufficient to fill a 300 mL box to a concentration of 5 ul/l (EthylBloc, AgroFresh). We then tightly sealed the lids of the boxes with parafilm. For cytokinin treatment experiments, 150 uM stock *trans-*zeatin in .01 N KOH was added to autoclaved media to create 150 nM working concentration (Sigma, Z0876). For the NPA experiment, 50 mM stock NPA in DMSO was first diluted in sterile water and added to autoclaved media to create a final concentration of 40 nM (Sigma, 33371). An equivalent amount of DMSO was used for the negative control. 200 mM IAA in ETOH was first diluted in sterile water and then added to hydroponically growing rice roots approximately 1 day after germination to create a final concentration of 200 nM (GoldBio, I-110-25) . 200 mM 1-NAA and 2-NAA stock solutions in acetone were diluted in sterile water and added to hydroponically grown rice roots approximately 1 day after germination to create a final concentration of 200 nM (Sigma, N0640 and N4002).

### RNA-Sequencing

Single 2mm root tip sections were isolated from 2 day post-germination seedlings. Sections were initially harvested into 10 ul of RNA-later (Ambion) in the lid of a 2 ml screw cap homogenization tube. 200 ul of Tri-Reagent (Zymo) was added to the tube, along with 2 3-mm stainless steel grinding balls, and the cap containing the tissue section was closed on the tube. It was inverted several time to wash the tissue section into the Tri-Reagent. Samples were frozen in liquid nitrogen and stored at -80, then processed by grinding with a bead homogenizer with 30 second pulses, 1500 hz, until the Tri-Reagent thawed (3-4 cycles). RNA was isolated using the Zymo MagBead RNA Isolation kit according to manufacturer’s protocol (Zymo). 100-200 ng of RNA was used as input into the Quantseq 3’ FWD RNA-Seq library preparation procedure according to the manufacturer’s protocol, using the Unique Molecular Identifier/UMI PCR addon kit (Lexogen). Libraries were indexed and pooled one lane of Illumina NextSeq, High Output setting. Reads were aligned to MSU Rice genome v7 using the STAR aligner (*44*), deduplicated using UMI-Tools (*45*), and counted with HTSeq-Count. Counts were analyzed with edgeR (*46*). RNA-Seq data analysis was performed by importing the counts output by HT-Seq into R for subsequent analysis with the EdgeR package ^8^. We defined “expressed genes” to be those with observed reads in 3 or more libraries. Files containing processed CPMs are included in Data S5 and) raw reads have been deposited at the Sequence Read Archive under BioProject ID PRJNA615381.

### Platform Exploration Assay

Seeds were sown in Magenta vessels containing media solidified with 0.5% Gelzan. Containers had raised polycarbonate surfaces of equal height with holes of 2.5mm diameter equally spaced in a square grid at distances of 5, 7, 9, and 11mm. Plants were grown on a robotic imaging system. We modified the robot by adding a second camera oriented at a 45 degree angle downward to visualize the growth of the roots on the platforms.

The containers were imaged for a minimum period of five days. A trial was considered a success when the primary root hit a surface and found a hole within 72 hours of first hitting the surface; failures occurred when the root hit the surface but did not find a hole in that time period. Exclusions included trials where the plant had no primary roots, the primary root never hit the surface or directly found the hole without surface contact, the primary root grew out of view of the camera, or the growth media was visibly contaminated.

### Statistical Analysis of Platform Exploration Assay

11, 12, 6, and 6 roots of wildtype and 12, 16, 18, and 7 roots of mutant were allowed to grow on a platform for 72 hours at hole spacings of 5mm, 7mm, 9mm, and 11mm respectively. “Success” was defined as the root tip encountering and growing into a hole. We modelled the probability of success using logistic regression with hole spacing and genotype as covariates. Based on the Wald test, we observed effects of genotype and spacing with p-values = 0.000211 and 0.007853, respectively. We utilized Wald based Confidence Interval estimation to determine confidence intervals for the odds ratios (95% CI for Odds Ratio for Genotype: (3.43, 45.09), 95% CI for spacing: (0.4966817, 0.8628481)). An analysis script, along with raw data, has been included in Data S6.

### Robotic Root Design

To generate 2D oscillation at the tip, we used two series pneumatic artificial muscles (sPAMs (*37*, *38*)) attached to the left and right side of the main body (see Fig.5B). SPAMs include multiple pouch motors (*47*) that are made of lay-flat poly tubing (d_tube = 25 mm) with O-rings (d_ring = 4.5 mm) spaced at 2 cm intervals. When the sPAMs are inflated, the segments separated by O-rings extend radially and shorten longitudinally. The shortening length of the sPAM bends the tip of the robot to the corresponding side. A key innovation is our method of localizing oscillation to only the tip. In (*37*), the entire body bends when an sPAM is actuated. To achieve tip-only oscillation, we incorporate a second, internal growing body inside the main growing outer body. This internal body grows at the same rate as the outer body, but is shorter and at a higher pressure. In this manner, the tip of the robot is formed by the outer body only, and is thus at a lower pressure than the rest of the body. Therefore, when an sPAM is actuated, only the lower pressure tip bends.

### Robotic Root Peg Assay

The robotic root was tested in an obstacle course to examine the exploratory capabilities of the nutation behavior. Peg obstacles were placed in a triangular lattice as described in (*38*). A transparent plastic sheet kept the robot root constrained to the 2-dimensional plane of obstacles.

### Definition of Robot Root Contact

To measure the effectiveness of obstacle traversal, we analyzed what occurred after the robot tip first hit an obstacle. The touch of the robot tip was considered “hitting” if the angle (α) between the line that connects the tip to the center of a peg and the line passing from the center of a peg to the center of the root approximately 15 cm away is less than approximately 60° (Fig. S8). Fig. S8 shows two snapshots from the experiment where the left panel shows an example of “hitting” and the right panel shows an example of contact not labelled as “hitting.”

### Peg Interaction Measurements

We performed two sets of experiments (21 experiments per each case) on the regular obstacle lattice; with a period of 5s and without tip oscillation starting from different initial conditions. The horizontal starting position of the robot was changed in ± 6 cm increments up to ± 25 distance from the center of the upper edge of the lattice. At each initial condition we performed three experiments. We tracked the robot tip from the video frames (1 frames/s) using MATLAB.

We counted the total number of times each root hit an obstacle before either becoming stuck or reaching the maximum extrusion length of about 1 meter. To estimate an overall metric of obstacle traversal effectiveness, we counted the total number of times any root hit an obstacle and successfully grew beyond. We divided this number by the total number times any root hit an obstacle.

### Peg Angle Measurements

We analyzed each interaction of the root with a peg and measure the angle of the root, as defined above, and whether the interaction resulted in a traversal or immobilization, for both nutating and non-nutating roots.

### Statistical Analysis of Peg Interaction

A Chi-Squared test for independence was performed on the 2x2 contingency table corresponding to the successes and failures of the nutating and non-nutating robotic root in growing past an obstacle. An analysis script, along with raw data, has been included in Data S7.

### Gravel Exploration Assay

Seeds were surface sterilized and pregerminated for 1 day in sterile liquid media at 30 dC in the dark before being surface sown onto a 1.5 cm layer of liquid saturated autoclaved calcined clay gravel (Turface MVP) on top of a 100 ml volume of .15% gelzan solidified Yoshida’s media inside magenta boxes, 9 seeds total per box. These boxes were imaged in the automated robotic imaging system for 4 days. We defined 1 day post-sowing as germination, and then counted the number of roots emerging from the gravel layer into the gel layer 3 days later. This quantity gave the numerator for our calculation of the proportion of success. We calculated the denominator by counting the number of seeds that had germinated with an emerged primary root. We planted a total of 11 boxes of both wildtype and mutant, with a total of 88 and 71 germinating wildtype and mutant seeds, respectively.

### Statistical Analysis of Gravel Exploration

A Chi-Squared test for independence was performed on the 2x2 contingency table corresponding to the successes and failures of the wildtype and *hk1-2* roots growing into the gel layer 3 days after germination. An analysis script, along with raw data, has been included in Data S8.

### Video Processing

For plant growth, time-lapse movies were created with ffmpeg at a framerate of 15 frames/second (FFmpeg Project), except for the platform experiment where the video was created in Matlab at 10 frames/second. Since images were taken every 15 minutes, this translates to 225 minutes growth/1 second of video (or 150 minutes growth/second in the platform video). Videos were stabilized and brightness/contrast adjusted with Premiere Pro 2019 (Adobe). In some videos masks were applied along the edge of the video to occlude roots from adjacent plants growing into the field of view. In other cases masks were applied to portions of the field of view unconnected to the seed or root where blemishes in the box created small regions of scattered light. Aside from stabilization and brightness/contrast adjustment, the region of the movie occupied by the seed or root was not modified.

**Fig. S1).**
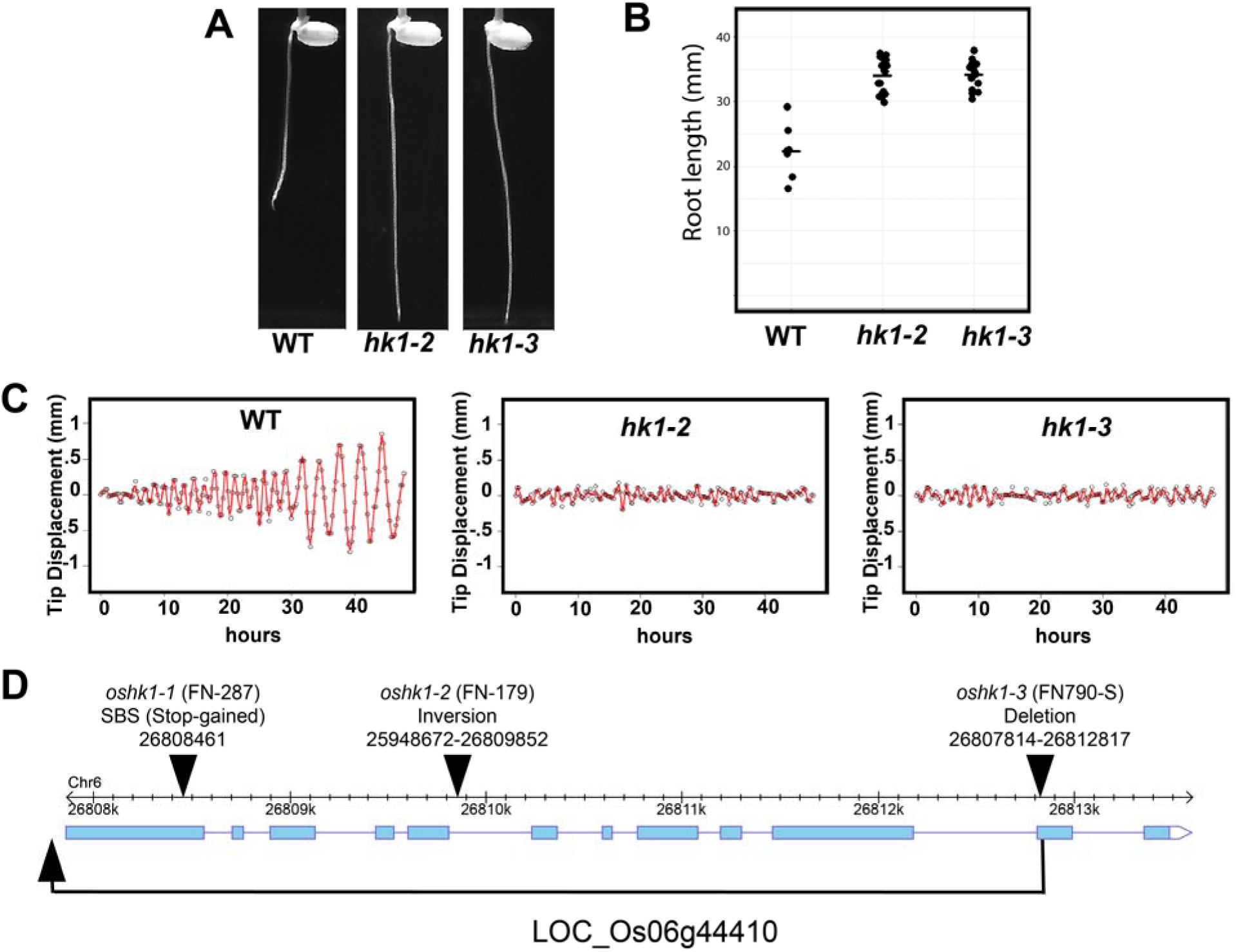
Identification of allelic mutants of *hk1* with identical deeper primary root depth and circumnutation phenotypes. A) Primary root growth of wildtype, *hk1-2,* and *hk1-3* 48 hours after germination B) Quantification of primary root length comparing wildtype to *hk1-2* and *hk1-3.* C) Tip traces of representative roots for wildtype, *hk1-2,* and *hk1-3* over 48 hours of growth D) Diagram of mutations in *hk1-1, hk1-2,* and *hk1-3.*

**Fig. S2).**
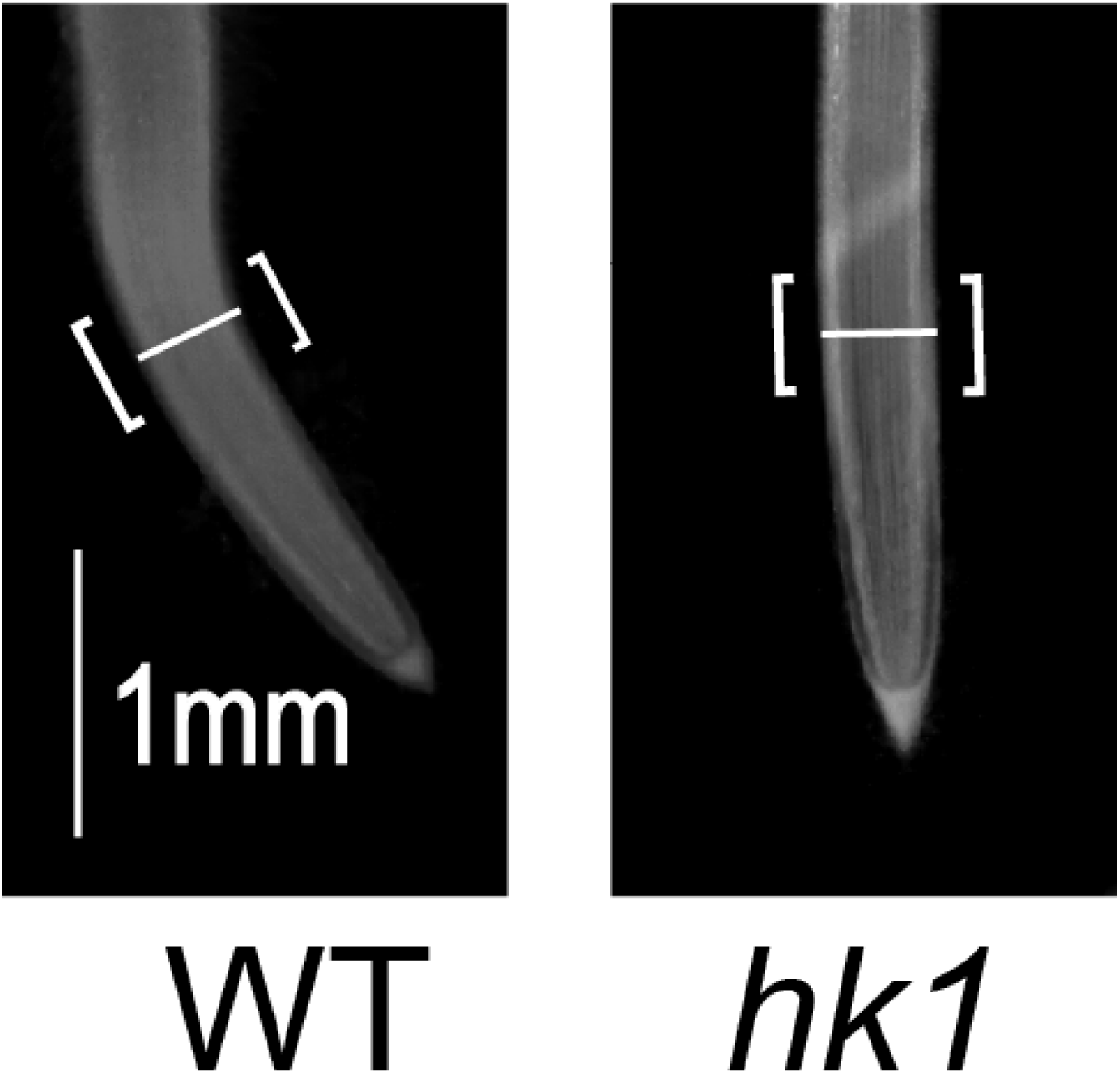
Region of cell length measurements in wildtype and mutant.

**Fig. S3).**
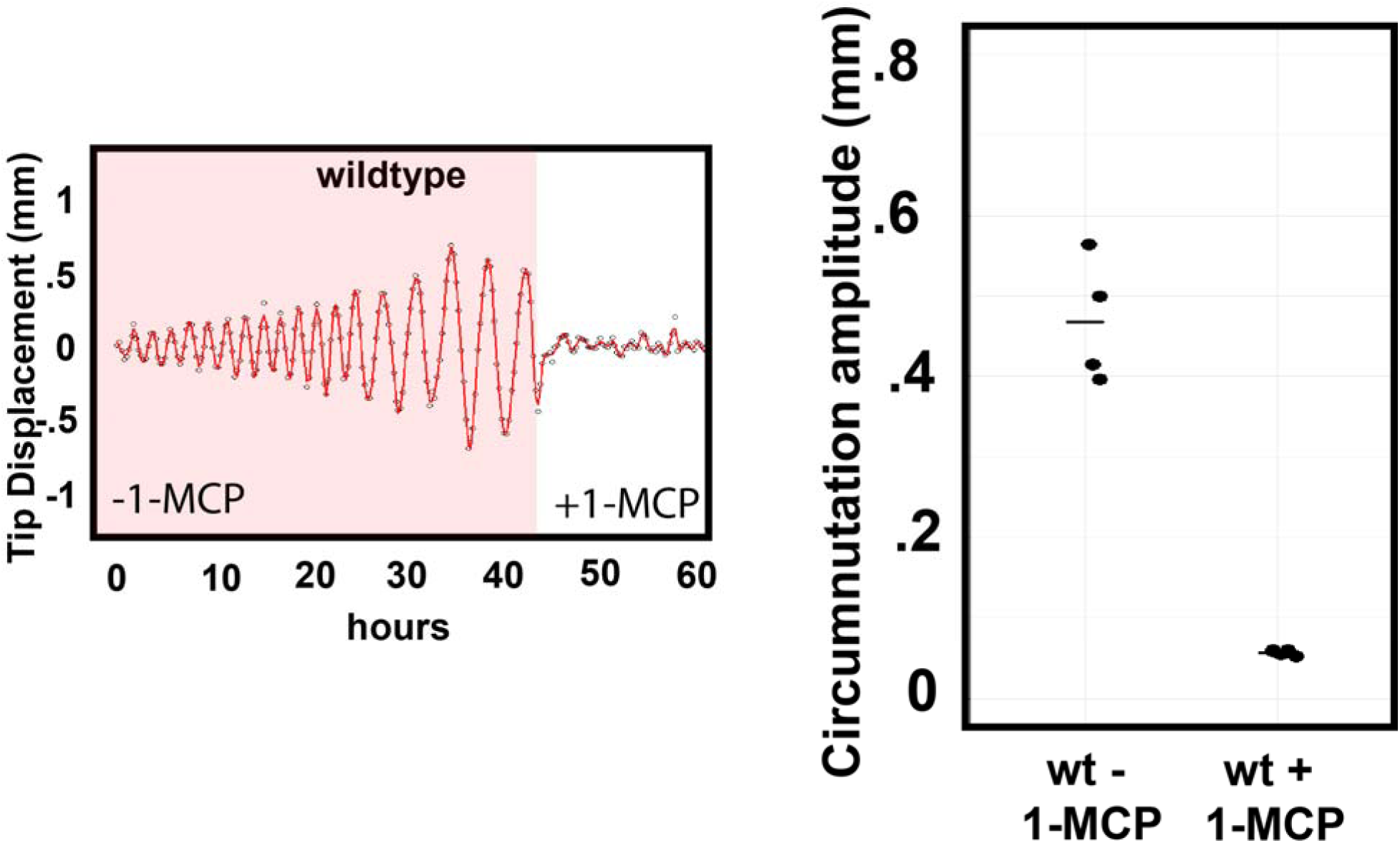
Treatment of wildtype with the ethylene receptor inhibitor 1-MCP blocks circumnutation. Graph displays average amplitude of circumnutation after time-point of 1-MCP addition for 4 and 5 untreated and treated roots, respectively. Two-sided Wilcoxon rank-sum test p-value = 0.01587. Horizontal dash indicates mean value.

**Fig. S4).**
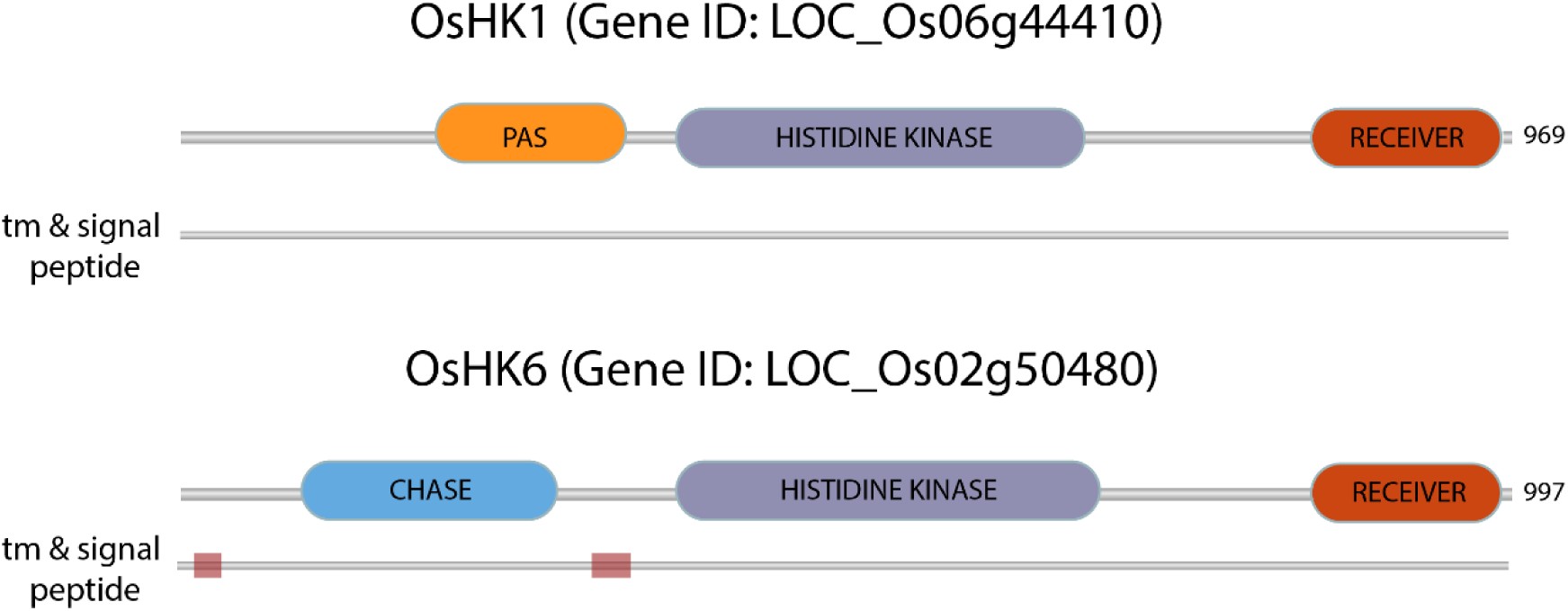
Domain architecture of HK1 and a predicted canonical cytokinin receptor, HK6.

**Fig. S5).**
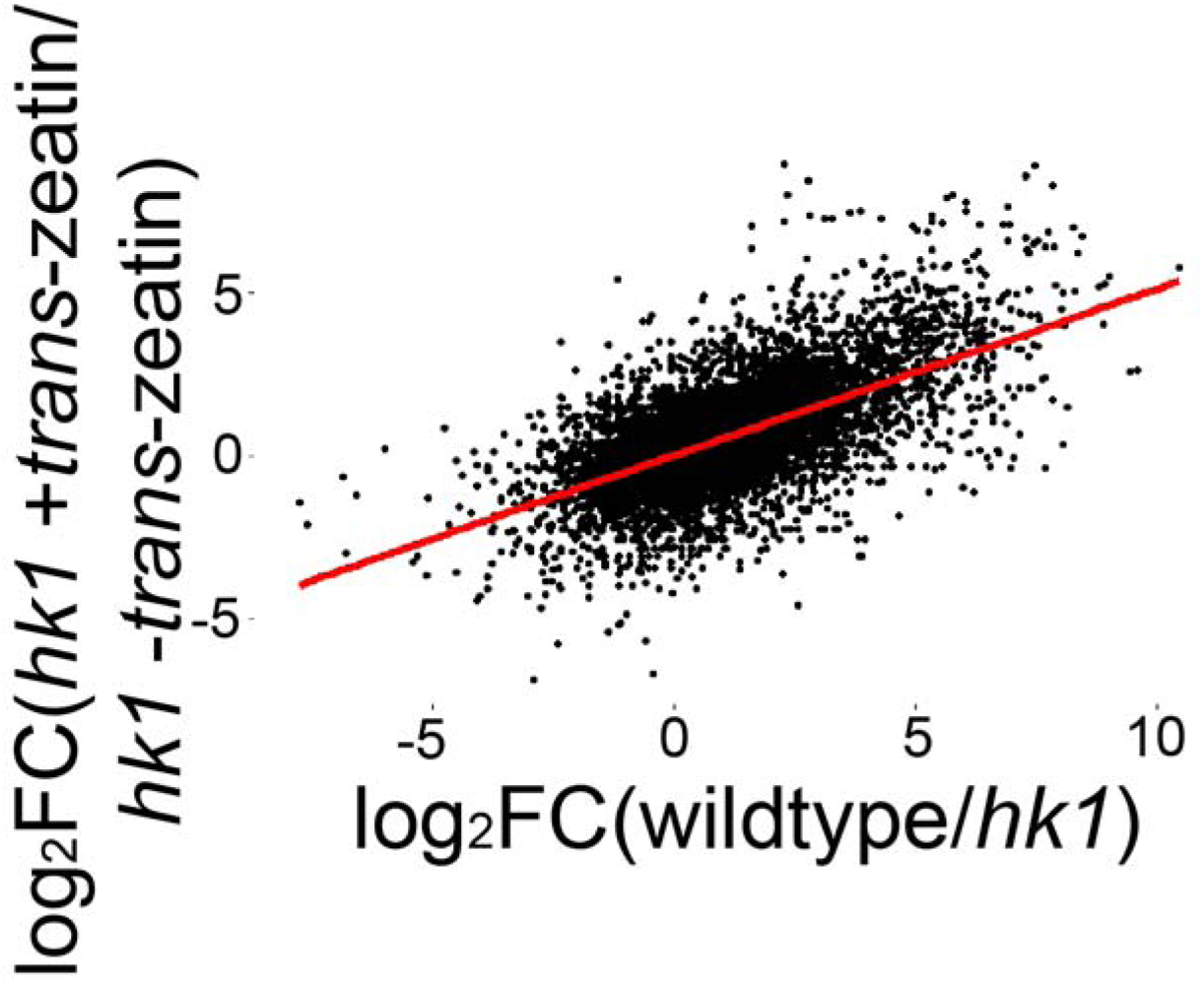
log2(fold change) values for all genes in wildtype/*hk1* compared with *hk1* plus/minus *trans-*zeatin

**Fig. S6).**
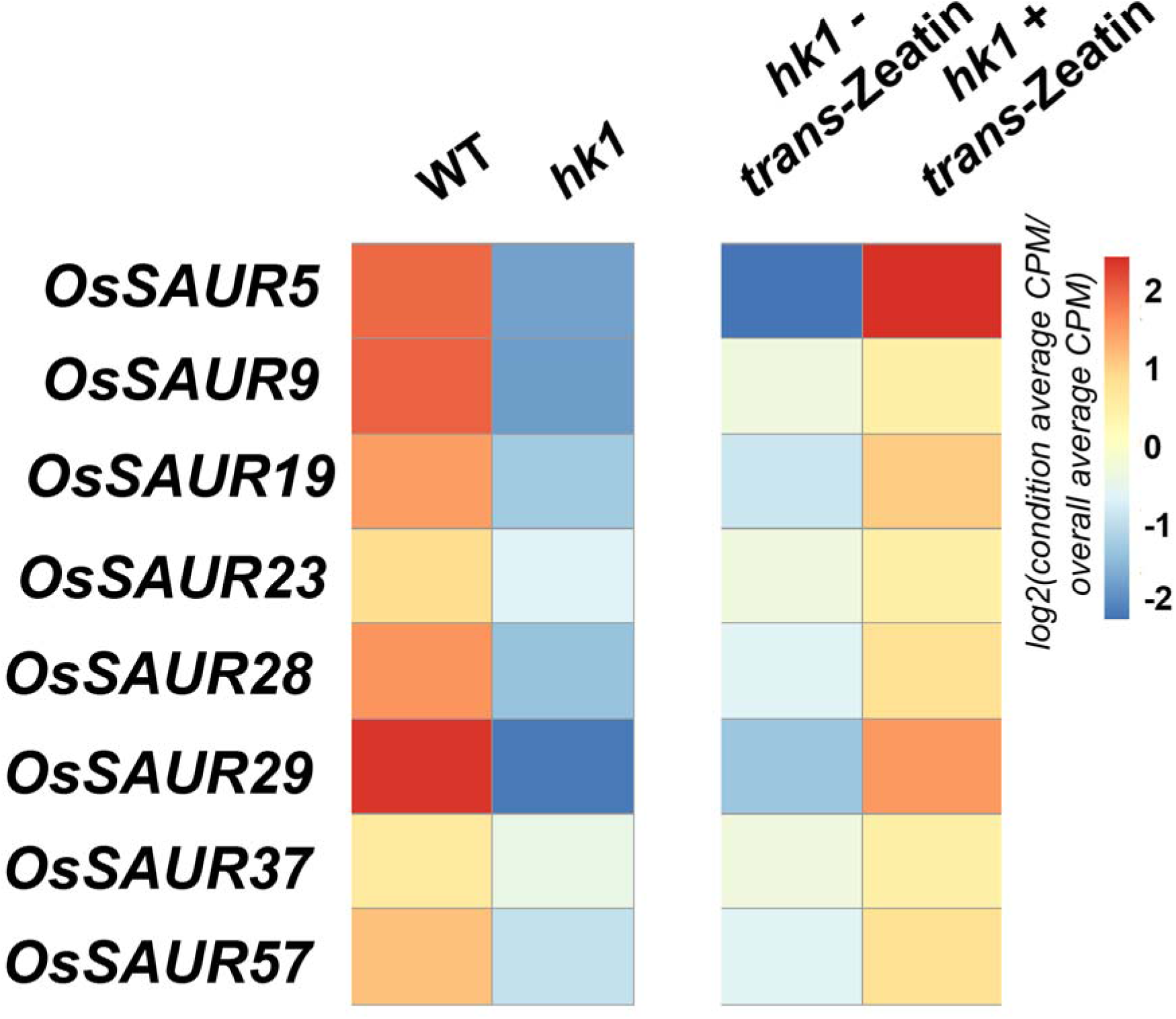
log2(fold change) of 8 differentially expressed *SAUR* genes for wildtype/*hk1* and *hk1* plus/minus *trans-*zeatin

**Fig. S7).**
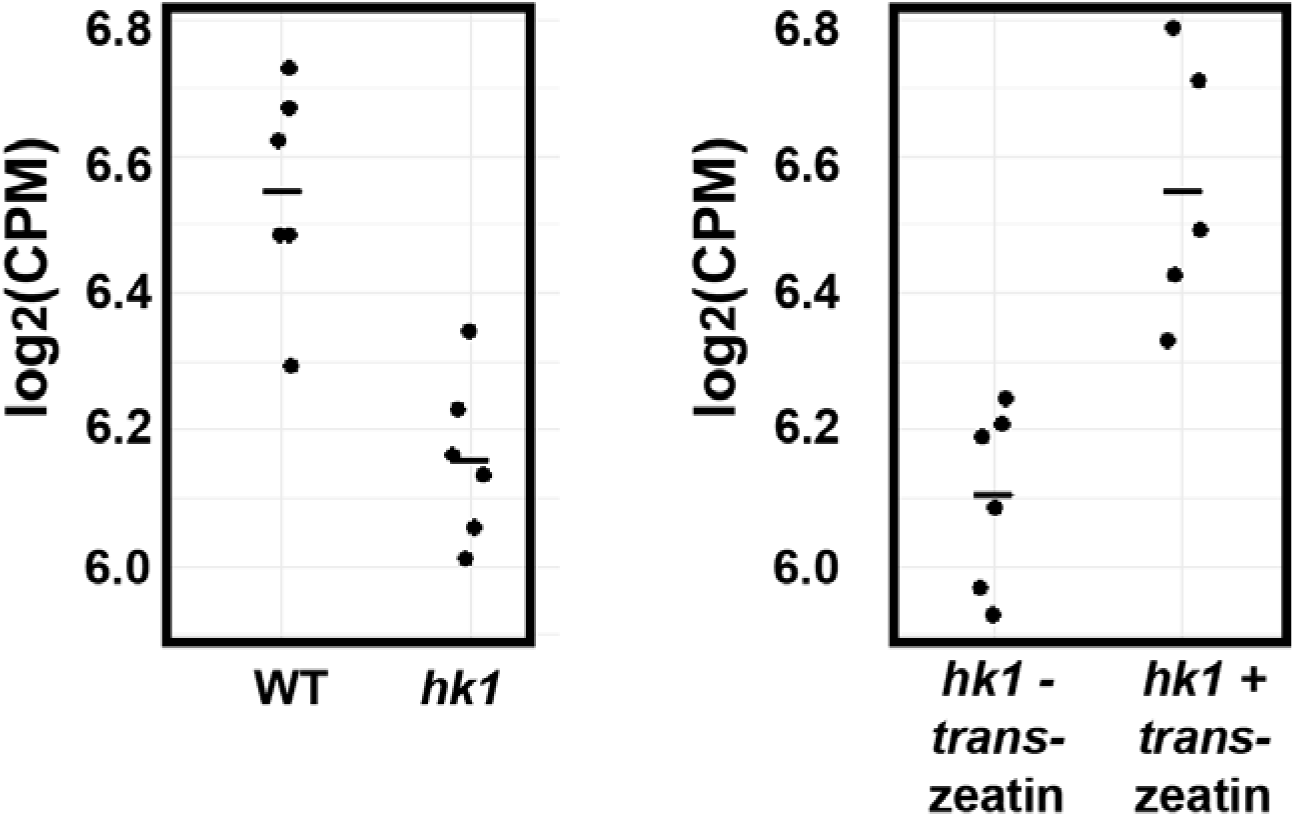
log2(counts per million/CPM) for *AUX1* in wildtype vs *hk1* and *hk1* plus/minus *trans*-zeatin

**Fig. S8).**
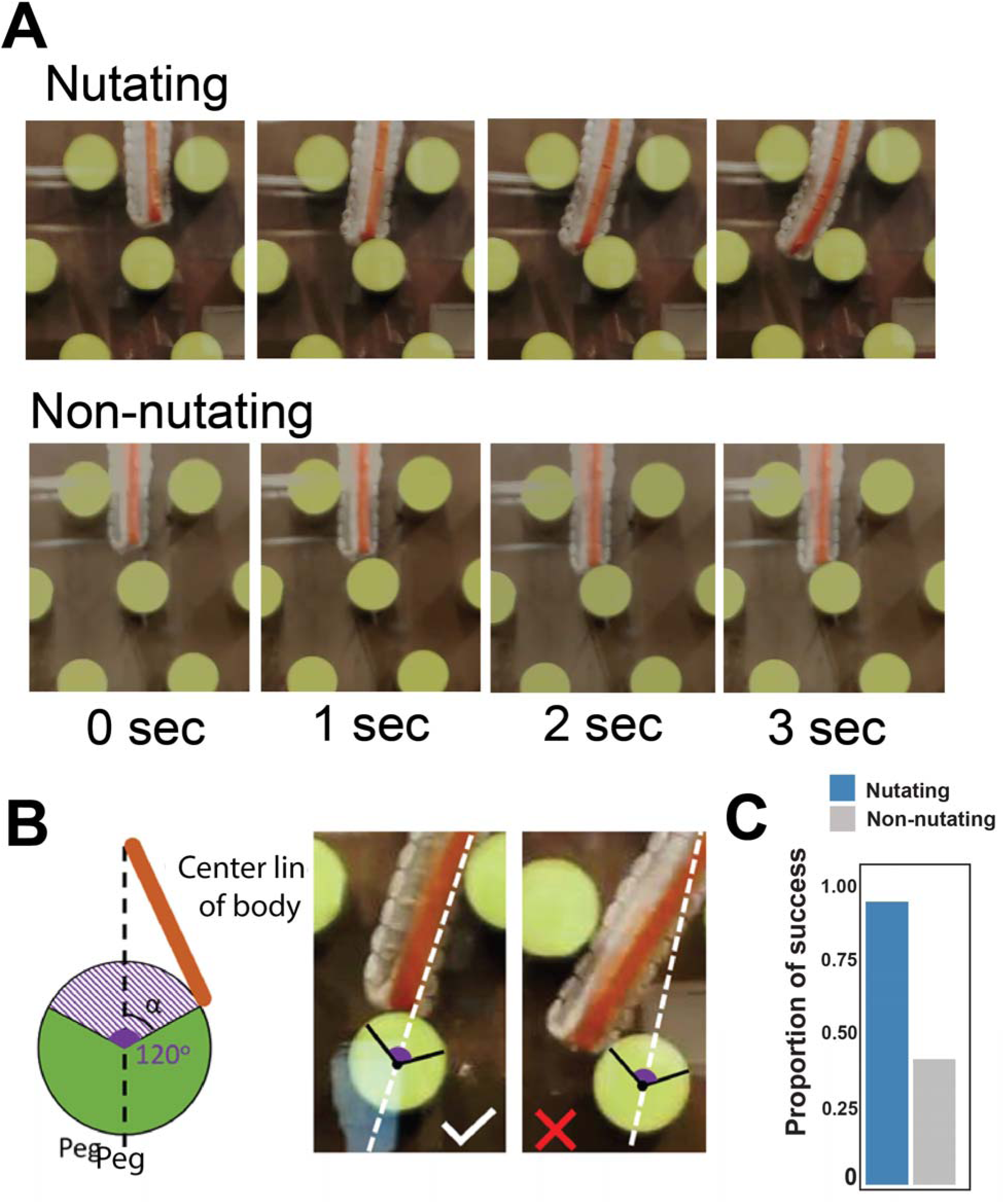
Definition of an “encounter” during robotic root growth. A) Example of nutating and non-nutating roots striking the same peg with a similar angle. B) The touch of the robot tip is counted as an “encounter” if the angle (α) between the line that connects the tip to the center of a peg and the line passing from the center of a peg is less than approximately 60 degrees. The left panel shows an example of an “encounter” and the right panel does not. C) The overall proportion of successful obstacle clearances. 21 experiments in both conditions, with a total of N = 59 and 33 obstacles encountered for nutating and nonnutating, respectively. Chi-Squared test p-value = 7.768e-10.

**Fig. S9).**
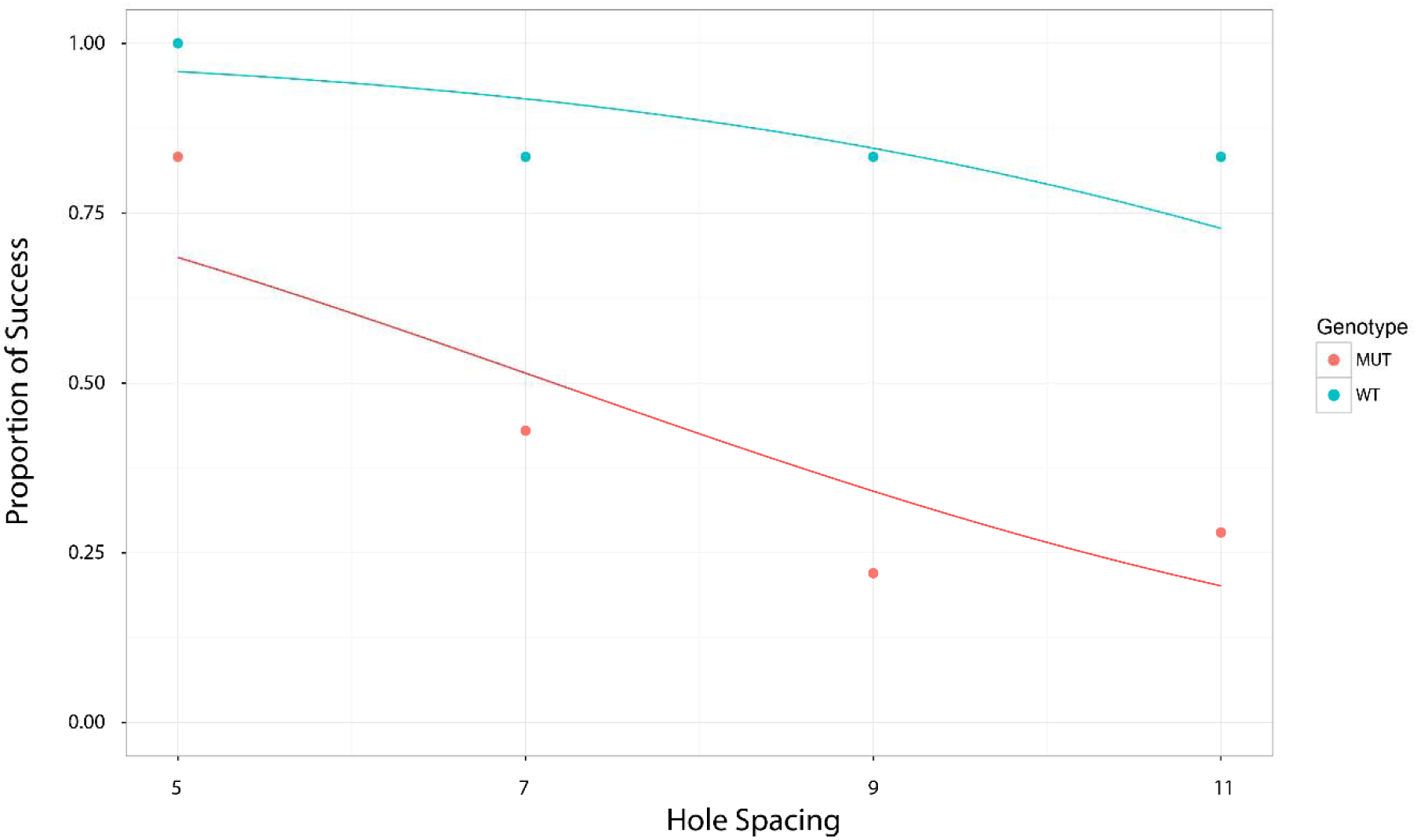
Proportion of success in finding hole for wildtype and mutant along with fitted logistic regression functions.

## Movie S1

Circumnutation phenotypes of wildtype and hk1-2. Video depicts 105 hours of growth.

## Movie S2

Circumnutation phenotype of wildtype before and after treatment with 1-MCP. Video depicts 86 hours of growth. 1-MCP applied at 13 second mark.

## Movie S3

Circumnutation phenotypes of untreated *hk1-2* and *hk1-2* treated with 150 nM *trans-*zeatin. Video depicts 90 hours of growth.

## Movie S4

Circumnutation phenotypes of untreated wildtype and wildtype treated with 40 nM NPA. Video depicts 64 hours of growth.

## Movie S5

Circumnutation phenotypes of hydroponically grown *hk1-1* treated with 200 nM IAA, 200 nM 2-NAA, and 200 nM 1-NAA. Video depicts 41 hours of growth. Treatments were applied at the 4 second mark.

## Movie S6

Circumnutation phenotypes of wildtype (Dongjin cultivar) and the *aux1-3* mutant. Video represents 94 hours of growth.

## Movie S7

Surface growth phenotypes of wildtype and *hk1-1*. Video depicts 63 hours of growth.

## Movie S8

Example circumnutating and non-circumnutating robotic simulation of root growth through an obstacle course of irregularly spaced pegs.

## Movie S9

Germination phenotype of wildtype and *hk1-2* on clay gravel. Video depicts 109 hours of growth.

## Movie S10

Germination phenotype of wildtype and *hk1-2* on 1.5 cm layer of clay gravel overlaying a layer of clear gel. Video depicts 135 hours of growth.

## Data S1

Compressed .Rproj file containing analysis script and raw data for primary root length measurement. Open .Rproj file and run “main” script in “scripts” folder.

## Data S2

Compressed .Rproj file containing script and raw data for root tip trace plotting. Open .Rproj file and run “main” script in “scripts” folder.

## Data S3

Compressed .Rproj file containing analysis script and raw data for circumnutation amplitude quantification. Open .Rproj file and run “main” script in “scripts” folder.

## Data S4

Compressed .Rproj file containing analysis script and raw data for epidermal cell length measurement. Open .Rproj file and run “main” script in “scripts” folder.

## Data S5

Processed count files for RNA-Seq experiments.

## Data S6

Compressed .Rproj file containing analysis script and raw data surface exploration experiment. Open .Rproj file and run “main” script in “scripts” folder.

## Data S7

Compressed .Rproj file containing analysis script and raw data robotic root experiment. Open .Rproj file and run “main” script in “scripts” folder.

## Data S8

Compressed .Rproj file containing analysis script and raw data for gravel penetration experiment. Open .Rproj file and run “main” script in “scripts” folder.

## Notes

### Competing Interest Statement

The authors have declared no competing interest.

