## Supplementary material for "Mechanism and function of root circumnutation": Data S1: main.R

```
#THIS SCRIPT READS IN LENGTH DATA FOR ROOTS OF WILDTYPE AND THE 3 hk1 ALLELES, PLOTS THEM, AND CALCULATES THE 
#WILCOXON RANK SUM TEST STATISTIC FOR THE WT/hk1 COMPARISONS
#DATA IS READ IN FROM THE "wt_mutant_length_position.csv" SPREADSHEET IN THE data DIRECTORY IN THE RPrOJECT FOLDER.

library(ggplot2)
library(here)

mpp=10/83.435  #mm per pixel for this experiment

#read in data
my_data=read.csv(here("data", "wt_mutant_length_position.csv"), header=T)
my_data$length = my_data$root_length_px*mpp


#plot hk1-1.WT controls are from boxes 267 and 268.
ggplot(my_data[(my_data[["genotype"]]=="wt" & as.integer(my_data[["plant"]])<269) | my_data[["genotype"]]=="hk1_1",], aes(x=genotype, y=length, group = genotype)) +
  scale_x_discrete(limits=c("wt", "hk1_1"))+ 
  geom_jitter(width = 0.05, size = 3) +
  ylim(0,42) +
  theme_bw() +
  stat_summary(fun.y= mean, fun.ymin=mean, fun.ymax=mean, geom="crossbar", width=0.2, color="black")

#wilcox test wt vs hk1-1
wilcox.test(my_data[my_data[["genotype"]]=="wt" & as.integer(my_data[["plant"]])<269, ]$root_length_px, my_data[my_data[["genotype"]]=="hk1_1",]$root_length_px)

#plot hk1-2 and hk1-3. WT controls are from boxes 269 and 270

ggplot(my_data[(my_data[["genotype"]]=="wt" & as.integer(my_data[["plant"]])>268) | my_data[["genotype"]]=="hk1_2"| my_data[["genotype"]]=="hk1_3",], aes(x=genotype, y=length, group = genotype)) +
  scale_x_discrete(limits=c("wt","hk1_2","hk1_3"))+ 
  geom_jitter(width = 0.05, size = 3) +
  ylim(0,42) +
  theme_bw() +
  stat_summary(fun.y= mean, fun.ymin=mean, fun.ymax=mean, geom="crossbar", width=0.2, color="black")


#wilcox.text wt vs hk1-2
wilcox.test(my_data[my_data[["genotype"]]=="wt" & as.integer(my_data[["plant"]])>268, ]$root_length_px, my_data[my_data[["genotype"]]=="hk1_2",]$root_length_px)

#wilcox.text wt vs hk1-3
wilcox.test(my_data[my_data[["genotype"]]=="wt" & as.integer(my_data[["plant"]])>268, ]$root_length_px, my_data[my_data[["genotype"]]=="hk1_3",]$root_length_px)
```
