## Supplementary material for "Mechanism and function of root circumnutation": Data S2: loess_spline_functions.R

```
#function to plot curves and calculate amplitudes. coords is the dataframe of coordinates. start and stop are datapoints being plotted.
#mpp is the pixel conversion. spn is the span for the loess model. It is increased for the 1-NAA experiment because root growth was
#extremely slow.
plot_curve <- function (coords, start, stop, mpp, spn){
  xy_df=coords[start:stop,2:3] #columns 2 and 3 are the coordinates, only take rows we're interested in plotting
  
  #loess regression to estimate center line of root. span = .2 for almost everything, .6 for 1-NAA treated
  loessMod10 <- loess(BX~BY, data=xy_df, span=spn) #use Loess to estimate centerline. Span can be tuned. BX and BY are from imageJ
  plot(BX~BY, data=xy_df)
  lines(xy_df$BY, predict(loessMod10, data.frame(BY = xy_df$BY)))
  
  
  #correct for drift
  BX_corrected=0 #will be the BX adjusted to the point along the loess prediction surface nearest to the observed point
  BX_min=0 #keeps track of shortest distance
  dist=0 #placeholder variable for distance
  
  #count is a counter variable
  count=0
  for (obs in xy_df$BY){
    count=count+1 #track which observation is being analyzed
    BX_min=xy_df[count,1]-predict(loessMod10, xy_df[count,])
    for (offset in c(-6,-5,-4,-3,-2,-1,0,1,2,3,4,5,6)) { #the pixel offset range we're scanning over. Can tune.
      if(!is.na(predict(loessMod10, obs+offset))) #deal with predicting at the ends where offsets will produce NAs
        dist=euc.dist(xy_df[count,],c(predict(loessMod10, obs+offset),obs+offset)) #calculate distance of each point to point along curve with offset
      #print(dist)
      if(dist<abs(BX_min)) {
        BX_min=BX_min/abs(BX_min)*dist #save offset to minimum if it's smaller than the previously found minimum. Preserve sign.
      }
    }
    BX_corrected[count]=BX_min
  }
  
  #plot individual points
  plot(BX_corrected*mpp~c(seq(start-1,(stop-1)/4,.25)), ylim=c(-1.25,1.25), xlab="hours",ylab="distance from center line (millimeters)")
  #plot a spline over the corrected values
  spline.lm <- lm((BX_corrected*mpp) ~ ns(seq(start-1,(stop-1)/4,.25), df=100), data=xy_df)
  lines(seq(start-1,(stop-1)/4,.25), predict(spline.lm), lwd=2, col='red')
}

#euclidean distance function. 
euc.dist <- function(x1, x2) sqrt(sum((x1 - x2) ^ 2))


#local maximum function. From user stas g. https://stackoverflow.com/questions/34205515/finding-local-maxima-and-minima-in-r
find_peaks <- function (x, m = 3){
  shape <- diff(sign(diff(x, na.pad = FALSE)))
  pks <- sapply(which(shape < 0), FUN = function(i){
    z <- i - m + 1
    z <- ifelse(z > 0, z, 1)
    w <- i + m + 1
    w <- ifelse(w < length(x), w, length(x))
    if(all(x[c(z : i, (i + 2) : w)] <= x[i + 1])) return(i + 1) else return(numeric(0))
  })
  pks <- unlist(pks)
  pks
}
```
