## Supplementary material for "Mechanism and function of root circumnutation": Data S2: main.Rmd

````
---
title: "loess_spline circumnutation R Notebook"
output: html_notebook
#THIS NOTEBOOK GENERATES THE PLOTS OF TIP DISPLACEMENT SHOWN IN THE PAPER. THE COORDINATES ARE READ IN FROM THE "all_tip_tracking_combined.csv" FILE IN THE data DIRECTORY IN THE RPROJECT FOLDER. THE TIMEPOINTS ARE SEPARATED BY 15 MINUTES
---

```{r}
library(ggplot2)
library(splines)
library(here)
source(here("functions", "loess_spline_functions.R")) #function needed for plotting. They are located in the "functions" subdirectory in the RPROJECT
```

```{r}
#estimate sequence specific distance factor 
mpp1=10/83.435  #mm per pixel conversion for camera 1
mpp2=10/244  #mm per pixel conversion for camera 2
```

```{r}
#read data. "plant" is the experiment number and the specific plant. "BX"" and "BY"" are the tip coordinates, "range" are the frames after germination recorded (1_192 means frames 1-192, and since each frame represents 1/4 of an hour, 192*1/4=48 hours of measurements). uid is a unique identifier corresponding to the plant and the range, "line" is the experiment number.
all_data=read.csv(here("data","all_tip_tracking_combined.csv"), header=T)
all_data[,5]=paste(all_data[,1],all_data[,4])
all_data[,6]=as.integer(all_data[,1])
colnames(all_data)=c("plant","BX","BY", "range", "uid", "line")
```


Make plots
```{r}
#wt mutant
plot_curve(all_data[all_data$uid=="267.1 1_192",], 1, 192, mpp1, .2) #WT
plot_curve(all_data[all_data$uid=="282.1 1_192",], 1, 192, mpp1, .2) #mutant

#NPA. 
plot_curve(all_data[all_data$uid=="884.4 1_192",], 1, 192, mpp1, .2) #plus NPA
plot_curve(all_data[all_data$uid=="890.1 1_192",], 1, 192, mpp1, .2) #minus NPA

#aux1
plot_curve(all_data[all_data$uid=="414.2 1_192",], 1, 192, mpp1, .2) #Dongjin
plot_curve(all_data[all_data$uid=="598.1 1_192",], 1, 192, mpp1, .2) #aux1

#tZ
plot_curve(all_data[all_data$uid=="592.1 1_233",], 1, 233, mpp1, .2) #plus tZ
plot_curve(all_data[all_data$uid=="594.2 1_233",], 1, 233, mpp1, .2) #minus tZ

#1-NAA
plot_curve(all_data[all_data$uid=="600.1 m48_p95",], 1, 144, mpp1, .2) #minus 1-NAA
plot_curve(all_data[all_data$uid=="602.2 m48_p95",], 1, 144, mpp1, .4) #plus 1-NAA. Use larger spanin LOESS. Like .4

#1-MCP
plot_curve(all_data[all_data$uid=="1091.3 germ-parafilm+10hr",], 1, 250, mpp2, .2) #minus 1-MCP
plot_curve(all_data[all_data$uid=="1095.1 germ-parafilm+10hr",], 1, 250, mpp2, .2) #plus 1-MCP

```
````
