## Supplementary material for "Mechanism and function of root circumnutation": Data S3: main.Rmd

````
---
title: "amplitude calculation"
output: html_notebook
#THIS SCRIPT UTILIZES A MODIFIED VERSION OF THE LOESS/SPLINE SCRIPT TO CALCULATE THE AMPLITUDE OF CIRCUMNUTATION IN A DEFINED WINDOW OF TIME (THE WINDOWS IS CHOSEN TO INCLUDE SEVERAL OSCILLATIONS OF THE TIP, AND THE AMPLITUDE IS ESTIMATED AS THE MEAN OF THESE OSCILLATIONS). DATA IS READ IN FROM THE SAME FILE "all_tip_tracking_combined.csv" LOCATED IN THE data SUBDIRECTORY OF THIS RPROJECT
---

```{r}
library(ggplot2)
library(splines)
library(here)
source(here("functions", "amplidtude_calculations_functions.R")) #function called in this script, located in "functions" subdirectory of Rproject folder
```

```{r}
#estimate sequence specific distance factor 
mpp1=10/83.435  #mm per pixel conversion for camera 1
mpp2=10/244  #mm per pixel conversion for camera 2 (near the end of our study we upgraded to a higher res camera, and a few experiments were performed with it)

```

```{r}
#read data in. "plant" is the experiment number and the specific plant. "BX"" and "BY"" are the tip coordinates, "range" are the frames after germination recorded (1_192 means frames 1-192, and since each frame represents 1/4 of an hour, 192*1/4=48 hours of measurements). uid is a unique identifier corresponding to the plant and the range, "line" is the experiment number.
all_data=read.csv(here("data","all_tip_tracking_combined.csv"), header=T)
all_data[,5]=paste(all_data[,1],all_data[,4])
all_data[,6]=as.integer(all_data[,1])
colnames(all_data)=c("plant","BX","BY", "range", "uid", "line")
```


```{r}

#WT VS hk1 alleles. FIRST 864 OBSERVATIONS ARE FOR THIS EXPERIMENT. FIG 1
#make data.frame for data
sub_data=all_data[1:864,]
average=data.frame(matrix(, ncol=2, nrow=27))
colnames(average)=c("line","av_displacement")
average[["line"]]=as.integer(unique(sub_data[,1]))
average[["genotype"]]=c(rep("wt",7), rep("it30",7), rep("it31", 6), rep("it32",7))

count=0
for(j in unique(sub_data[["plant"]])) {
  count=count+1
  peaks=(plot_curve2(sub_data[sub_data[["plant"]]==j,], 1, 32, mpp1, 100))
  indices= sort(peaks[34:(length(peaks))])
  avg=mean(abs(peaks[indices[2:(length(indices)-1)]]))
  average[count,2]=avg
}

average[["line"]]=as.factor(average[["line"]])
average[["genotype"]]=as.factor(average[["genotype"]])


#plot WT vs hk1.  it32= hk1-1 it30 = hk1-2. it31 = hk1-3
ggplot(average[1:27,], aes(x = genotype, y = av_displacement)) +
  scale_x_discrete(limits=c("wt", "it32","it30","it31"))+ 
  ylim(c(0,.8)) +
  geom_jitter(width=.1, size = 3) +
  theme_bw()+
  stat_summary(fun.y = mean, fun.ymin=mean, fun.ymax=mean, geom="crossbar", width=.2, color="black")

#CALCULATE MEDIANS AND WILCOXON RANK SUM TEST P-VALUES
median(average[average$genotype=="wt",]$av_displacement) #median wildtype amplitude = 0.563147
median(average[average$genotype=="it32",]$av_displacement) #median mutant amplitude = 0.1307679
wilcox.test(average[average$genotype=="wt",]$av_displacement,average[average$genotype=="it32",]$av_displacement) #p-value = 0.0005828


#hk1 minus vs hk1 + cytokinin. 591 and 592 are plus, 593 and 594 are minus
#make data.frame for data
sub_data=all_data[all_data$line %in% c(591, 592, 593, 594) & all_data$range == "201_233",]
average=data.frame(matrix(, ncol=2, nrow=length(unique(sub_data$plant))))
colnames(average)=c("line","av_displacement")
average[["line"]]=as.integer(unique(sub_data[,1]))
average[["treatment"]]=c(rep("mut+",2), rep("mut-",3), rep("mut-", 2), rep("mut+",2))

count=0
for(j in unique(sub_data[["plant"]])) {
  count=count+1
  peaks=(plot_curve2(sub_data[sub_data[["plant"]]==j,], 1, 32, mpp1, 100))
  indices= sort(peaks[34:(length(peaks))])
  avg=mean(abs(peaks[indices[2:(length(indices)-1)]]))
  average[count,2]=avg
}

average[["line"]]=as.factor(average[["line"]])
average[["treatment"]]=as.factor(average[["treatment"]])


#plot hk1 plus and minus cytokinin. 
ggplot(average, aes(x = treatment, y = av_displacement)) +
  scale_x_discrete(limits=c("mut-", "mut+"))+ 
  ylim(c(0,.8)) +
  geom_jitter(width=.1, size = 3) +
  theme_bw()+
  stat_summary(fun.y = mean, fun.ymin=mean, fun.ymax=mean, geom="crossbar", width=.2, color="black")


median(average[average$treatment=="mut-",]$av_displacement) #median tZ untreated amplitude = 0.09767216
median(average[average$treatment=="mut+",]$av_displacement) #median tZ treated amplitude = 0.6773349
wilcox.test(average[average$treatment=="mut-",]$av_displacement, average[average$treatment=="mut+",]$av_displacement) # p-value = 0.01587

#wildtype with DMSO or NPA. 40 nM. 
sub_data=all_data[all_data$line %in% c(884, 885, 890, 891) & all_data$range == "160_192",]
average=data.frame(matrix(, ncol=2, nrow=length(unique(sub_data$plant))))
colnames(average)=c("line","av_displacement")
average[["line"]]=as.integer(unique(sub_data[,1]))
average[["treatment"]]=c(rep("WT+",5), rep("WT-",6))

count=0
for(j in unique(sub_data[["plant"]])) {
  count=count+1
  peaks=(plot_curve2(sub_data[sub_data[["plant"]]==j,], 1, 33, mpp1, 100))
  indices= sort(peaks[34:(length(peaks))])
  avg=mean(abs(peaks[indices[2:(length(indices)-1)]]))
  average[count,2]=avg
}

#plot WT DMSO vs NPA
ggplot(average, aes(x = treatment, y = av_displacement)) +
  scale_x_discrete(limits=c("WT-", "WT+"))+ 
  ylim(c(0,.8)) +
  geom_jitter(width=.1, size = 4) +
  theme_bw()+
  stat_summary(fun.y = mean, fun.ymin=mean, fun.ymax=mean, geom="crossbar", width=.2, color="black")

median(average[average$treatment=="WT-",]$av_displacement) #median NPA untreated amplitude = 0.6438134
median(average[average$treatment=="WT+",]$av_displacement) #median NPA treated amplitude = 0.07310549
wilcox.test(average[average$treatment=="WT-",]$av_displacement, average[average$treatment=="WT+",]$av_displacement) # p-value = 0.004329

#hk1 - and + NAA
sub_data=all_data[all_data$line %in% c(599, 600, 601, 602) & all_data$range == "m48_p95",]
average=data.frame(matrix(, ncol=2, nrow=length(unique(sub_data$plant))))
colnames(average)=c("line","av_displacement")
average[["line"]]=as.integer(unique(sub_data[,1]))
average[["treatment"]]=c(rep("mut-",7), rep("mut+",7))

count=0
for(j in unique(sub_data[["plant"]])) {
  count=count+1
  peaks=(plot_curve2(sub_data[sub_data[["plant"]]==j,], 1, 144, mpp1, 100))
  indices= sort(peaks[(length(peaks)-4):(length(peaks))])
  avg=mean(abs(peaks[indices[2:(length(indices)-1)]]))
  average[count,2]=avg
}

#plot hk1 and hk1 + 1-NAA
ggplot(average, aes(x = treatment, y = av_displacement)) +
  scale_x_discrete(limits=c("mut-", "mut+"))+ 
  ylim(c(0,.9)) +
  geom_jitter(width=.1, size = 3) +
  theme_bw()+
  stat_summary(fun.y = mean, fun.ymin=mean, fun.ymax=mean, geom="crossbar", width=.2, color="black")

median(average[average$treatment=="mut-",]$av_displacement) #median 1_NAA untreated amplitude = 0.1663442
median(average[average$treatment=="mut+",]$av_displacement) #median 1-NAA treated amplitude = 0.6675895
wilcox.test(average[average$treatment=="mut-",]$av_displacement, average[average$treatment=="mut+",]$av_displacement) # p-value = 0.0005828


#Dongjin vs aux1
sub_data=all_data[all_data$line %in% c(415, 597, 598) & all_data$range == "161_192",]
average=data.frame(matrix(, ncol=2, nrow=length(unique(sub_data$plant))))
colnames(average)=c("line","av_displacement")
average[["line"]]=as.integer(unique(sub_data[,1]))
average[["treatment"]]=c(rep("WT",5), rep("mut",5))

count=0
for(j in unique(sub_data[["plant"]])) {
  count=count+1
  peaks=(plot_curve2(sub_data[sub_data[["plant"]]==j,], 1, 32, mpp1, 100))
  indices= sort(peaks[(length(peaks)-1):(length(peaks))])
  avg=mean(abs(peaks[indices[2:(length(indices)-1)]]))
  average[count,2]=avg
}

#plot Dongjin vs aux1
ggplot(average, aes(x = treatment, y = av_displacement)) +
  scale_x_discrete(limits=c("WT", "mut"))+ 
  ylim(c(0,.5)) +
  geom_jitter(width=.1, size = 3) +
  theme_bw()+
  stat_summary(fun.y = mean, fun.ymin=mean, fun.ymax=mean, geom="crossbar", width=.2, color="black")

median(average[average$treatment=="WT",]$av_displacement) #median Dongjin WT amplitude = 0.2863864
median(average[average$treatment=="mut",]$av_displacement) #median aux1-3 amplitude = 0.08017139
wilcox.test(average[average$treatment=="WT",]$av_displacement,average[average$treatment=="mut",]$av_displacement) # p-value = 0.007937

#MCP +/-
sub_data=all_data[all_data$line %in% c(1090, 1091, 1094, 1095) & all_data$range == "parafilm+2hr-8hr",]
average=data.frame(matrix(, ncol=2, nrow=length(unique(sub_data$plant))))
colnames(average)=c("line","av_displacement")
average[["line"]]=as.integer(unique(sub_data[,1]))
average[["treatment"]]=c(rep("minus",4), rep("plus",5))

count=0
for(j in unique(sub_data[["plant"]])) {
  count=count+1
  peaks=(plot_curve2(sub_data[sub_data[["plant"]]==j,], 1, 34, mpp2, 25))
  indices= sort(peaks[(length(peaks)-1):(length(peaks))])
  avg=mean(abs(peaks[indices[2:(length(indices)-1)]]))
  average[count,2]=avg
}

#plot MCP
ggplot(average, aes(x = treatment, y = av_displacement)) +
  scale_x_discrete(limits=c("minus", "plus"))+ 
  ylim(c(0,.8)) +
  geom_jitter(width=.1, size = 4) +
  theme_bw()+
  stat_summary(fun.y = mean, fun.ymin=mean, fun.ymax=mean, geom="crossbar", width=.2, color="black")

median(average[average$treatment=="minus",]$av_displacement) #median 1-MCP untreated WT amplitude = 0.4568151
median(average[average$treatment=="plus",]$av_displacement) #media 1-MCP treated WT amplitude = 0.05740331
wilcox.test(average[average$treatment=="minus",]$av_displacement, average[average$treatment=="plus",]$av_displacement) # p-value = 0.01587

```
````
