## Supplementary material for "Mechanism and function of root circumnutation": Data S4: main.R

```
#THIS SCRIPT READS IN THE MEASURED CELL LENGTHS ON THE OUTER AND INNER BENDS OF WILDTYPE AND "SIDE 1" AND "SIDE 2" IN MUTANT

library(readxl)
library(ggplot2)
library(here)

#intialize dataframe
wt_data=data.frame(matrix(, nrow=0, ncol=6))
colnames(wt_data) = c("midline_distance","length", "side", "root", "angle", "angle_start")

mut_data=data.frame(matrix(, nrow=0, ncol=6))
colnames(mut_data) = c("midline_distance","length", "side", "root", "angle_start")

#vectors for differences
wt_diff=c()
mut_diff=c()

for(i in 2:10) { #loop through sheets of excel notebook
  #We create a dataframe with the following columns:
  #midline_distance = distance of the cell along the middle path of the root where the cell is located (um)
  #length  = length of the cell (um)
  #side = inner or outer
  #root = root, numbered 1-9
  #angle = the measured angle of the root
  #angle_start = the distance along the midline that is considered the maximal bend.
  
  #read in data
  wt_temp <- read_excel(here("data","Analysis_WT_final_graphs.xlsx"), sheet = i)
  #subset portion of interest
  wt_data1 = data.frame(wt_temp[7:dim(wt_temp)[1],2:4])
  #convert to double
  wt_data1[,1:2]=lapply((wt_data1[,1:2]), as.double)
  #add col for root
  wt_data1 = cbind(wt_data1, rep(i-1, dim(wt_data1)[1]))
  #add angle
  wt_data1$angle = rep(wt_temp[1,2], dim(wt_data1[1])[1])
  wt_data1$angle_start = as.double(rep(wt_temp[2,2], dim(wt_data1[1])[1]))
  #change colnames
  colnames(wt_data1) = c("midline_distance","length", "side", "root", "angle", "angle_start")
  

  #Here we identify which cell is located at the point of maximum bending (ie, is closest to the distance of maximum bending 
  #along the midline of the root). We do this for the inner and outer sides.
  inner_index=which(abs(wt_data1$midline_distance-wt_data1$angle_start[1])==min(abs(wt_data1[wt_data1$side=="Inner",]$midline_distance-wt_data1$angle_start[1])))
  outer_index=which(abs(wt_data1$midline_distance-wt_data1$angle_start[1])==min(abs(wt_data1[wt_data1$side=="Outer",]$midline_distance-wt_data1$angle_start[1])))
  
  #We sum up the 15-cell length by adding the all the lengths of all the cells 8 rootward, 6 shootward, and the cell at the
  #maximum bend itself. 
  wt_diff[i-1]=sum(wt_data1[(inner_index-8):(inner_index+6),]$length)-sum(wt_data1[(outer_index-8):(outer_index+6),]$length)

  #append data
  wt_data = rbind(wt_data, wt_data1)

}


for(i in 2:12) {#loop through sheets of excel notebook
  #We create a dataframe with the following columns:
  #midline_distance = distance of the cell along the middle path of the root where the cell is located (um)
  #length  = length of the cell (um)
  #side = arbitrarily labelled 0 or 1
  #root = root, numbered 1-11
  #angle = the angle of the root
  #angle_start, the distance along the midline that is a similar distance from the root tip as in wildtype
  
  #read in data
  mut_temp <- read_excel(here("data","Analysis_MUT_final_graphs.xlsx"), sheet = i)
  #get part that we want
  mut_data1 = data.frame(mut_temp[7:dim(mut_temp)[1],2:4])
  #convert to double
  mut_data1[,1:2]=lapply((mut_data1[,1:2]), as.double)
  #add col for root
  mut_data1 = cbind(mut_data1, rep(i-1, dim(mut_data1)[1]))
  #add angle
  mut_data1$angle = rep(mut_temp[1,2], dim(mut_data1[1])[1])
  mut_data1$angle_start = as.double(rep(mut_temp[2,2], dim(mut_data1[1])[1]))
  #change colnames
  colnames(mut_data1) = c("midline_distance","length", "side", "root", "angle", "angle_start")
  
  #Here we identify which cell is located at the point comparable to that of maximum bending (ie, is closest to the 
  #point along the midline of the root). We do this for the inner and outer sides.
  inner_index=which(abs(mut_data1$midline_distance-mean(mut_data1$midline_distance))==min(abs(mut_data1[mut_data1$side==1,]$midline_distance-mean(mut_data1$midline_distance))))
  outer_index=which(abs(mut_data1$midline_distance-mean(mut_data1$midline_distance))==min(abs(mut_data1[mut_data1$side==0,]$midline_distance-mean(mut_data1$midline_distance))))
  
  #We sum up the 15-cell length by adding the all the lengths of all the cells 8 rootward, 6 shootward, and the cell at the
  #point comparable to the maximum bend in the mutant. 
  mut_diff[i-1]=sum(mut_data1[(inner_index-8):(inner_index+6),]$length)-sum(mut_data1[(outer_index-8):(outer_index+6),]$length)

  #append data
  mut_data = rbind(mut_data, mut_data1)
}

#make categorical variable
wt_data$root = as.factor(wt_data$root)
wt_data$side = as.factor(wt_data$side)
mut_data$root = as.factor(mut_data$root)
mut_data$side = as.factor(mut_data$side)


#craft dataframe
dfwm=data.frame(matrix(, ncol=2, nrow=20))
dfwm[,1]=c(rep("wt",9), rep("mut",11))
dfwm[,2]=-1*c(wt_diff,mut_diff)
colnames(dfwm) = c("genotype","difference")

#plot differences
ggplot(dfwm, aes(x=genotype, y=difference, group = genotype)) +
  scale_x_discrete(limits=c("wt", "mut"))+ 
  geom_jitter(width = 0.05, size = 3) +
  ylim(min(dfwm[,2])-1, max(dfwm[,2])+1) +
  theme_bw() +
  stat_summary(fun.y= mean, fun.ymin=mean, fun.ymax=mean, geom="crossbar", width=0.2, color="black")

#test wt
wilcox.test(dfwm[dfwm$genotype=="wt",]$difference)

#test mutant
wilcox.test(dfwm[dfwm$genotype=="mut",]$difference)
```
