## Supplementary material for "Mechanism and function of root circumnutation": Data S6: main.R

```
library(ggplot2)
library(aod)
library(here)


#read in data
#3 columns
#Y_success = success or failure in growing through within 72 hours of striking the surface
#genotype= wildtype or hk1 mutant
#spacing= distance between holes in platform= 5, 7, 9, or 11 mm
y1=read.csv(here("data","logistic_data.csv"), header=T)

#create factor variable based on genotype
y1$Genotype = factor(y1$Genotype)

#fit logistic regression
mylogit <- glm(Y_success ~ Genotype + Spacing, data = y1, family = "binomial")

#summary of model with Wald tests
summary(mylogit)

#confidence intervals
exp(confint(mylogit)) #profile likelihood
exp(confint.default(mylogit)) #Wald

#graph regression function on data
WT_func=exp(predict(mylogit, newdata=data.frame("Spacing"=seq(5,11, .01), "Genotype"=factor("WT"), type="response")))

MUT_func=v2=exp(predict(mylogit, newdata=data.frame("Spacing"=seq(5,11, .01), "Genotype"=factor("MUT"), type="response")))
                
props=data.frame(rep(c(5,7,9,11), 2), c(1,.833,.833,.833,.833,.43,.22,.28),c(rep("WT",4), rep("MUT",4)))
colnames(props)=c("Spacing", "prop","genotype")
temp_WT=data.frame(seq(5,11,.01), WT_func/(1+WT_func),  rep("model_WT"))
colnames(temp_WT)=c("Spacing", "prop","genotype")
temp_MUT=data.frame(seq(5,11,.01), MUT_func/(1+MUT_func),  rep("model_MUT"))
colnames(temp_MUT)=c("Spacing", "prop","genotype")

#plot of logistic regression fit
f<-ggplot(props, aes(x=Spacing, y=prop))  +
  geom_point(aes(color=props$genotype), size=3) +
  scale_y_continuous(limits = c(0, 1)) +
  geom_line(data = temp_WT, aes(Spacing, prop), col="cyan3") + #COLOR
  geom_line(data = temp_MUT, aes(Spacing, prop), col="brown1") + #COLOR
  theme_bw() +
  labs(colour="Genotype", x="Spacing", y="proportion of success") +
  scale_x_continuous(breaks = seq(5, 11, by = 2)) 
f

props$genotype=factor(props$genotype, levels = c("WT","MUT"))
#barplot of success data
w<-ggplot(props, aes(x=Spacing, y=prop, fill=genotype))  +
  geom_bar(stat = "identity", position = "dodge") +
  labs(colour="Genotype", x="Spacing", y="proportion of success") +
  scale_x_continuous(breaks = seq(5, 11, by = 2)) +
  scale_fill_manual("legend", values = c("WT" =  "steelblue", "MUT" = "gray")) + #COLOR
  theme_bw() + theme(panel.grid.major = element_blank(),
                     panel.grid.minor = element_blank(), axis.line = element_line(colour = "black"))
w
```
