## Supplementary material for "Mechanism and function of root circumnutation": Data S7: main.Rmd

````
---
title: "robot nutation"
output: html_notebook
---

load packages
```{r}
library(ggplot2)
library(here)
library(survival)
library(survminer)
```

read data
```{r}
#angles daata
angles=read.csv(here("data","angles.csv"))
angles$abs=abs(angles$angle)
```


Overall proportion success
```{r}
#Calculate successful traversals
length(angles[angles$type=="OFF",]$success) #33
length(angles[angles$type=="ON",]$success) #59
sum(angles[angles$type=="OFF",]$success) #12
sum(angles[angles$type=="ON",]$success) #57


props_new = data.frame(matrix(ncol=2, nrow=2))
colnames(props_new)=c("Proportion_success","Condition")
props_new$Proportion_success=c(with_prop, without_prop)
props_new$Condition=c("WITH","WITHOUT")

props_new$Proportion_success=c(57/59, 12/33)

#barplot of Success data
w<-ggplot(props_new, aes(x=Condition, y=Proportion_success, fill=Condition))+  
  geom_bar(stat = "identity", position = "dodge") + 
  scale_y_continuous(limits= c(0,1))+
  labs(colour="Condition", x="Condition", y="Proportion_success")  +
  scale_fill_manual("legend", values = c("WITH" =  "steelblue", "WITHOUT" = "gray")) + #COLOR
  theme_bw() + theme(panel.grid.major = element_blank(),
                     panel.grid.minor = element_blank(), axis.line = element_line(colour = "black"))
w

#chi-squared test
prop.test(matrix(c(57,12,2,21),ncol=2)) #p-value = 7.768e-10
```


Success broken down by contact angle
```{r}
angles$bins= cut(angles$abs, breaks=c(0,10,20,30,70), labels = seq(10,40,10))


props=data.frame(matrix(ncol=3, nrow=8))
colnames(props)=c("type","bin","proportion")
props$type=c("OFF","OFF","OFF","OFF","ON","ON","ON","ON")
props$bin=c(10,20,30,40,10,20,30, 40)

props[1,3]=sum(angles[angles$type=="OFF" & angles$bins==10,]$success)/length(angles[angles$type=="OFF" & angles$bins==10,]$success)

props[2,3]=sum(angles[angles$type=="OFF" & angles$bins==20,]$success)/length(angles[angles$type=="OFF" & angles$bins==20,]$success)

props[3,3]=sum(angles[angles$type=="OFF" & angles$bins==30,]$success)/length(angles[angles$type=="OFF" & angles$bins==30,]$success)

props[4,3]=sum(angles[angles$type=="OFF" & angles$bins==40,]$success)/length(angles[angles$type=="OFF" & angles$bins==40,]$success)

props[5,3]=sum(angles[angles$type=="ON" & angles$bins==10,]$success)/length(angles[angles$type=="ON" & angles$bins==10,]$success)

props[6,3]=sum(angles[angles$type=="ON" & angles$bins==20,]$success)/length(angles[angles$type=="ON" & angles$bins==20,]$success)

props[7,3]=sum(angles[angles$type=="ON" & angles$bins==30,]$success)/length(angles[angles$type=="ON" & angles$bins==30,]$success)

props[8,3]=sum(angles[angles$type=="ON" & angles$bins==40,]$success)/length(angles[angles$type=="ON" & angles$bins==40,]$success)


w<-ggplot(props, aes(x=bin, y=proportion, fill=type))  +
  geom_bar(stat = "identity", position = "dodge") +
  labs(colour="Type", x="Angle", y="proportion of success") +
  scale_x_continuous(breaks = seq(10,60,10)) +
  scale_fill_manual("legend", values = c("ON" =  "steelblue", "OFF" = "gray")) + #COLOR
  theme_bw() + theme(panel.grid.major = element_blank(),
                     panel.grid.minor = element_blank(), axis.line = element_line(colour = "black"))
w
```
````
