## Supplementary material for "Mechanism and function of root circumnutation": Data S8: main.Rmd

````
---
title: "Gravel penetrations experiment"
output: html_notebook
#This script loads the data from "gravel.success.csv" in the "data" subdirectory of this Rproject, plots the proportion success in #roots growing into the gel layer 3 days after germination for widltype and mutant, and performs a chi-squared test for the #difference in proportion.
---

```{r}
library(here)
library(ggplot2)
```

```{r}
success <- read.csv(here("data", "gravel_success.csv"))
#Each box contained 9 seeds of the stated genotype. Then we counted the number of roots penetrating into the gel layer 3 days 
#after germination. We then excised all seedlings and counted the number that had germinated and had primary roots > 1.5 cm. In total these numbers form the numerator and denominator of the proportion of success. 
#are the numerator and denominator of 
#The plants were grown in 3 batches, 3 days post germination time for each batch is listed below.
#1351 WT 28s
#1352 WT 28s
#1353 WT 28s
#1354 WT 28s
#1355 WT 28s
#1356 hk1 28s
#1357 hk1 28s
#1358 hk1 28s
#1359 hk1 28s
#1360 hk1 28s

#1128 WT 23s
#1129 WT 23s
#1130 WT 23s
#1133 hk1 23s
#1134 hk1 23s
#1135 hk1 23s

#1180 WT
#1181 WT
#1182 WT
#1183 hk1
#1184 hk1
#1185 hk1

```

simple bar plot of success
```{r}
#make proportion
wt=sum(success[success$genotype=="wt",]$thru_72hr_pg)/sum(success[success$genotype=="wt",]$germinated)
hk1=sum(success[success$genotype=="hk1",]$thru_72hr_pg)/sum(success[success$genotype=="hk1",]$germinated)

props = data.frame(matrix(ncol=2, nrow=2))
colnames(props)=c("Proportion_success","Genotype")
props$Proportion_success=c(wt, hk1)
props$Genotype=c("WT","hk1")

#barplot of Success data
w<-ggplot(props, aes(x=reorder(Genotype, -Proportion_success), y=Proportion_success, fill=Genotype))+  
  geom_bar(stat = "identity", position = "dodge") + 
  scale_y_continuous(limits= c(0,1))+
  labs(colour="Genotype", x="Genotype", y="Proportion_success")  +
  scale_fill_manual("legend", values = c("WT" =  "steelblue", "hk1" = "gray")) + #COLOR
  theme_bw() + theme(panel.grid.major = element_blank(),
                     panel.grid.minor = element_blank(), axis.line = element_line(colour = "black"))
w
```

test
```{r}
sum(success[success$genotype=="wt",]$thru_72hr_pg)
sum(success[success$genotype=="hk1",]$thru_72hr_pg)
sum(success[success$genotype=="wt",]$germinated)
sum(success[success$genotype=="hk1",]$germinated)

tab=matrix(c(68,34,88-68,71-34), ncol=2)

prop.test(tab) #p-value = 0.000238
```
````
