## Supplementary figures and images for "Mechanism and function of root circumnutation"

### proportion_success.tiff

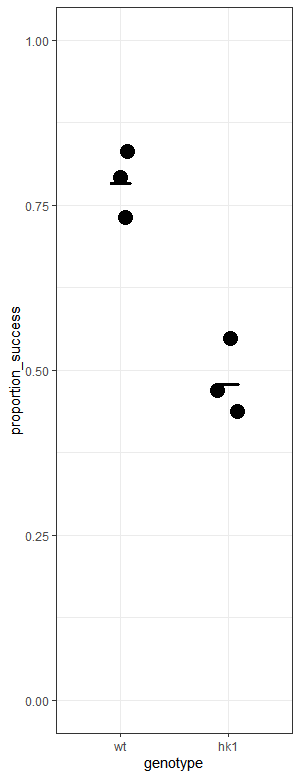
